# Reduced melanocortin tone mediates increased feeding during pregnancy in mice

**DOI:** 10.1101/2025.08.22.671837

**Authors:** Ingrid Camila Possa-Paranhos, Kerem Catalbas, Samuel Congdon, Tanya Pattnaik, Christina Nelson, Dajin Cho, Patrick Sweeney

**Author notes:** These authors contributed equally (Ingrid Camila Possa-Paranhos, Kerem Catalbas).

## Abstract

During pregnancy mammals increase their food intake to accommodate the elevated metabolic demands associated with fetal growth and development. However, the molecular and neural circuit mechanisms mediating increased feeding during pregnancy are largely unknown. Here, we demonstrate that arcuate nucleus agouti-related peptide (AgRP) neurons are activated and pro-opiomelanocortin (POMC) neurons are inhibited during pregnancy in mice. These changes are required for promoting hyperphagia during pregnancy as chemogenetic inhibition of AgRP neurons or activation of POMC neurons both reduced the feeding of pregnant mice to non-pregnant levels. Finally, we utilized single cell resolution spatial transcriptomics in the arcuate nucleus of non-pregnant and pregnant mice to characterize pregnancy-induced changes in the transcriptomic state of arcuate nucleus neurons, including significant changes in both AgRP and POMC neurons. Together, these findings outline a circuit mechanism mediating increased feeding during pregnancy, providing important mechanistic insights related to conditions at the intersection of reproduction and metabolism.

**Highlights:**

- AgRP neuron activity is increased and POMC neuron activity is decreased in pregnant mice
- Increased AgRP neuron activity and reduced POMC neuron activity is required for increased feeding during pregnancy
- Pregnancy enhances responsivity of AgRP neurons to palatable food
- Pregnancy drastically alters the transcriptional state of both AgRP and POMC neurons towards positive energy balance

## Introduction

Pregnancy presents an enormous metabolic challenge in mammals, requiring substantial increases in energy to promote fetal growth and development, and prepare for the subsequent lactational period. One prominent strategy employed by mammals to address these increased energy needs is to substantially increase food consumption.^1^ For example, rodents increase food intake by approximately 25% during the third trimester of pregnancy, while humans typically increase food intake by approximately 15% during this period in healthy pregnancies.^2, 3^ Although this increased feeding is essential to promote fetal growth and development, long-term maternal and fetal health is remarkably sensitive to maladaptive energy intake during pregnancy. Thus, in both humans and preclinical rodent models, overfeeding and underfeeding during pregnancy increases the risk of the mother and children developing metabolic disorders later in life.^4–6^ However, the core neural circuitry and molecular mechanisms mediating increased feeding during pregnancy are largely unknown.

Feeding behavior is primarily controlled by multiplexed neural circuitry in the hypothalamus and hindbrain, which initiate food seeking and consumption in response to reductions in energy availability and terminate feeding behavior in response to energy sufficiency.^7–9^ Decades of research supports an essential role for the hypothalamic melanocortin system in coordinating this process.^10, 11^ The melanocortin system is composed of agouti-related peptide (AgRP) and pro-opiomelanocortin (POMC) neurons in the arcuate nucleus, which exert antagonistic effects on feeding behavior and energy homeostasis. AgRP neurons are activated in response to energy deficiency to promote feeding and reduce energy expenditure, while POMC neurons are activated by energy sufficiency to reduce feeding and increase energy expenditure. AgRP neurons produce the endogenous melanocortin receptor inverse agonist/antagonist AgRP, while POMC neurons produce the endogenous melanocortin receptor agonist alpha-melanocyte stimulating hormone. Together, AgRP and aMSH regulate the activity of downstream melanocortin-4 receptors (MC4R) located in the paraventricular hypothalamus (PVN), with AgRP inhibiting MC4R activity to increase feeding and aMSH stimulating MC4R activity to reduce feeding.^10^ Gain of function mutations in AgRP and loss of function mutations in both POMC or MC4R result in hyperphagia and obesity in preclinical rodent models, fish, and humans.^12–18^ Further, loss of function mutations in MC4R are the most common monogenetic cause of obesity and hyperphagia in humans, occurring in nearly 1 in 300 individuals.^19, 20^ Thus, the central melanocortin pathway represents a remarkably well conserved pathway for controlling feeding behavior across a wide variety of species, including humans.

Although the melanocortin system is well established as a critical regulator of feeding behavior, the specific role of this pathway in regulating feeding during pregnancy remains enigmatic. Prior studies suggest that pregnant rats have increased levels of AgRP and reduced levels of POMC mRNA, although these findings are not consistent across studies.^21–23^ Consistently, pregnant rodents exhibit impairments in central responsivity to leptin and insulin, which together act on AgRP and POMC neurons to regulate feeding, energy homeostasis, and glycemia.^22^ The impairment of AgRP and leptin levels has also been observed in human cerebral spinal fluid or plasma samples, especially during the late pregnancy period.^24^ However, it is unclear if changes in the central melanocortin pathway are essential for promoting increased feeding during pregnancy, and the molecular mechanisms mediating increased feeding during pregnancy are largely unknown. Here, we utilized mouse feeding assays, fiber photometry, RNAscope in situ hybridization, chemogenetics, and spatial transcriptomics to characterize an essential role for the hypothalamic melanocortin system in promoting increased feeding during pregnancy in mice.

## Results

### Pregnancy increases feeding by increasing meal size in mice

Although pregnancy is known to increase feeding, the specific behavioral mechanism(s) mediating increased feeding during pregnancy are incompletely understood. Therefore, we first characterized food intake and body weight of pregnant and non-pregnant mice during pregnancy. Pregnant mice initially consumed similar amounts of food as non-pregnant mice during the early stages of pregnancy (**Fig. 1A**). However, starting in the mid-second to early third trimester of pregnancy, pregnant mice began to consume more food than non-pregnant mice, peaking at an approximately 25-30% increase in food intake during the third trimester of pregnancy (**Fig. 1A**). This increase in feeding was accompanied by a rapid increase in weight gain during the third trimester of pregnancy (**Fig. 1B**). To further measure feeding structure in undisturbed mice we utilized feeding experimental devices (FED3) to characterize feeding structure during the peak hyperphagic period of pregnancy (pregnancy day 15-19). FED3 devices attach directly to the mouse home-cage (**Fig. 1C**), allowing for the quantification of meal frequency and size in an undisturbed environment. Consistent with prior reports, pregnant mice consumed more food than non-pregnant mice (**Fig. 1D**), while consuming larger meals (**Fig. 1E**). Surprisingly, pregnant mice ate less frequently than non-pregnant mice (**Fig. 1F**). The distribution of meal sizes indicates that while most meals in non-pregnant mice consist of small meal sizes, pregnant mice exhibit a significant shift towards larger meals (**Fig. 1G**). Thus, pregnancy results in a shift in meal structure towards larger but more infrequent meals, perhaps facilitating increased feeding while conserving time and energy for other important physiological processes associated with pregnancy (i.e. nest building, reduced locomotion, etc).

**Figure 1:**
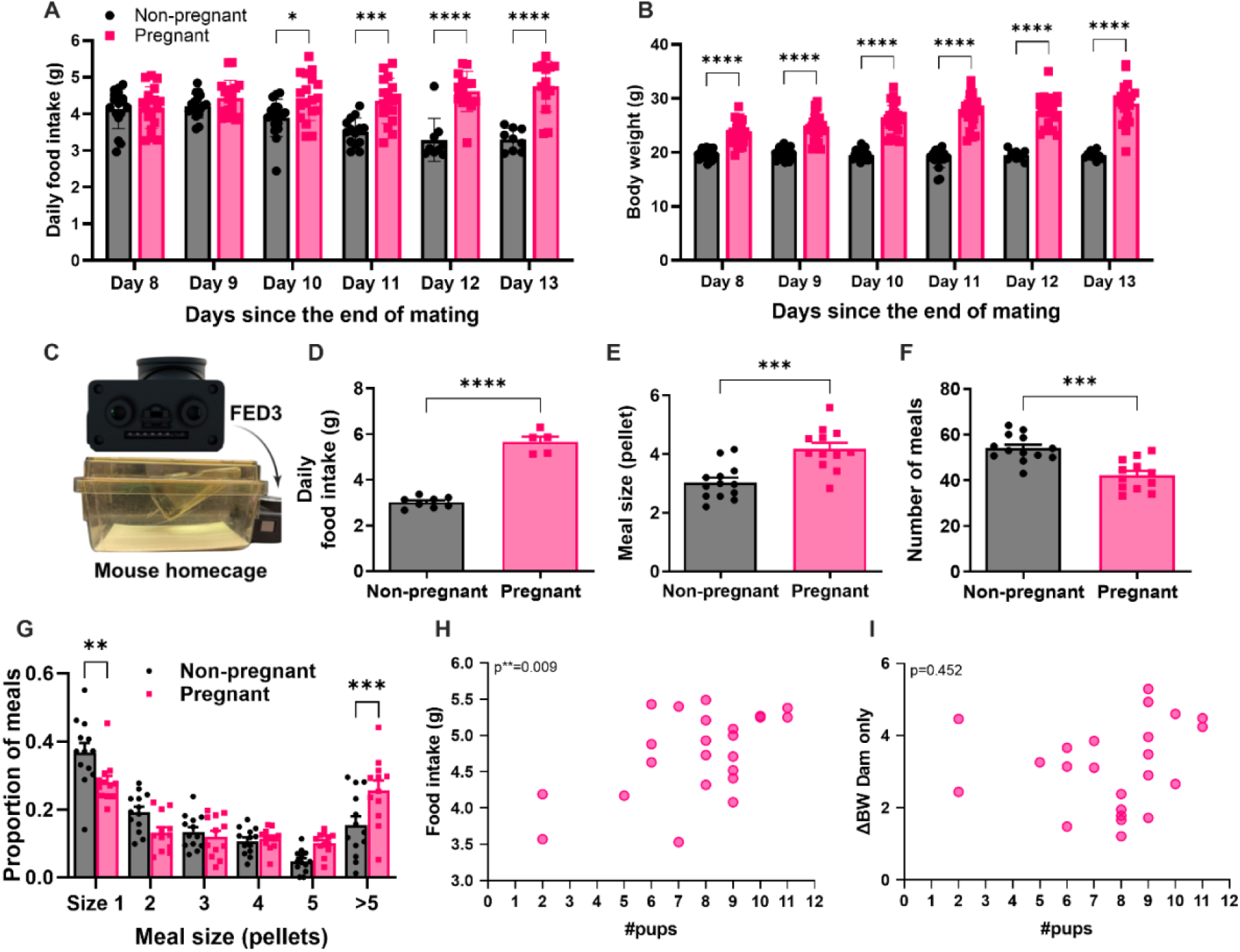
Hyperphagia present during pregnancy is mainly caused by increased meal size, not the number of meals. (A) Daily regular chow intake, in grams, in non-pregnant and pregnant animals from day 8 until day 13 after mating period (mice mated for 5 days). (n=10-18) (B) Comparison of daily body weight, in grams, in non-pregnant and pregnant animals from day 8 until day 13 after mating period. (n=10-18) (C) Set-up of the mouse home-cage with the FED3 feeding device. (D) Comparison between 24h food intake of non-pregnant and pregnant mice using the FED3 pellet dispenser. (n=5-8) (E) Number of pellets consumed per meal in 24h by non-pregnant and pregnant animals. (n=12-13) (F) Number of meals consumed in 24h by non-pregnant and pregnant mice. (n=12-13) (G) Comparison of the number of meals and their pellet size distribution in 24h. (n=12-13) (H) Correlation between regular chow 24h food intake of the pregnant mice and their number of pups. (n=23) (I) Correlation between regular chow 24h food intake of the pregnant mice and the body weight gain of the dam during the pregnancy period (subtracting the weight of the pups and the dam’s initial body weight before pregnancy) (n=23). Data are presented as mean values ± SEM. For panels A, B, and G, statistical significance was tested by 2way ANOVA and Šídák’s multiple comparisons test. For panels D-F, statistical significance was tested by Unpaired t test with Welch’s correction. For panels G-H, correlation was performed and tested by Pearson correlation coefficients. For all panels *p\** < 0.05, *p\*\** < 0.01, *p\*\** < 0.001. Data points represent individual mice.

To determine if the number of pups sired by the dam correlates with the body weight and food intake of the dam during pregnancy, following parturition we calculated the number of pups born and correlated this with the food intake and body weight gain of the dams during pregnancy. Food intake during pregnancy was positively correlated with the number of pups, with larger pregnancies resulting in increased levels of food intake (**Fig. 1H**). However, the total body weight gained by the dam (subtracting the weight of the pups and dam’s body weight prior to pregnancy to the dam’s body weight on the day of perfusion) did not correlate with the number of pups in gestation (**Fig. 1I**).

### Pregnancy increases mRNA expression of AgRP and reduces mRNA expression of POMC

After characterizing the microstructure of feeding in pregnant mice (**Fig. 1**), we next sought to identify the cell types mediating increased feeding during pregnancy. Given the established role for hypothalamic AgRP and POMC neurons in feeding behavior, we utilized RNAscope in situ hybridization to characterize AgRP and POMC mRNA levels in pregnant and non-pregnant mice (**Fig. 2A**). Consistent with the increased feeding behavior observed during pregnancy, pregnant mice had higher levels of AgRP mRNA than non-pregnant mice (**Fig. 2B, C, and E**). This increase in AgRP mRNA was particularly enhanced when normalizing the food intake levels between non-pregnant and pregnant animals (i.e. pair feeding pregnant mice to non-pregnant levels) (**Fig. 2D and E**). In contrast, mRNA levels of the anorexic melanocortin peptide POMC were significantly lower in pregnant mice, relative to non-pregnant control animals (**Fig. 2F, G, and I**). POMC peptide levels were further reduced when normalizing the level of food intake between pregnant and non-pregnant mice (i.e. by pair-feeding pregnant mice to non-pregnant levels of food intake; **Fig. 2H and I**). Thus, pregnancy concurrently increases mRNA levels of the orexigenic peptide AgRP and reduces mRNA levels of the anorexic peptide POMC.

**Figure 2:**
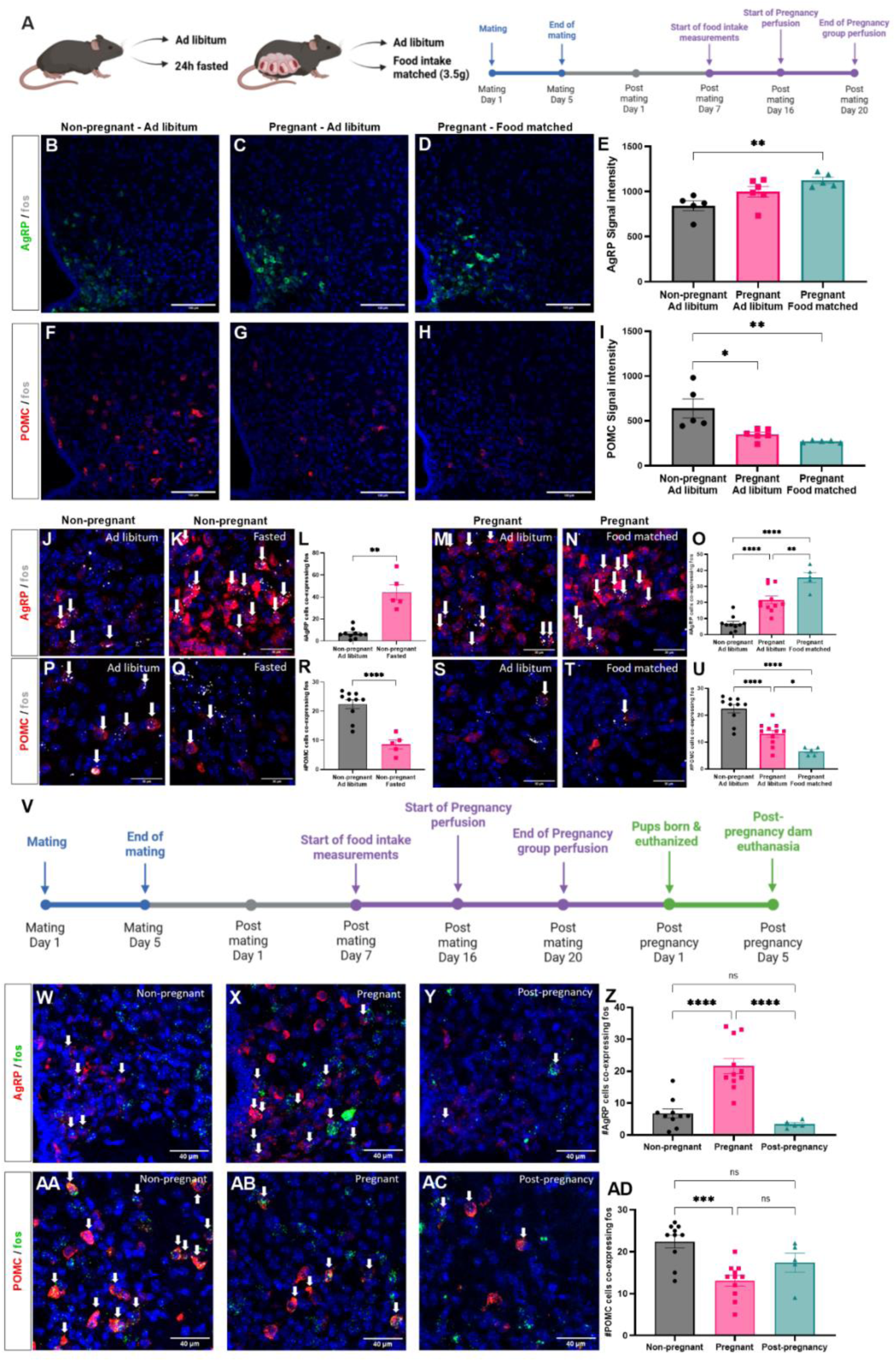
Pregnancy decreases POMC neuron activity and increases AgRP neuron activity. (A) Schematic and timeline of the pregnancy in situ hybridization experiment. (B-D) Confocal images of AgRP mRNA (green) expression and DAPI (blue) in Arc of female non-pregnant mouse with *ad libitum* access to food (B), pregnant female mouse with *ad libitum* access to food (C), and pregnant mouse with 24h hour food intake matched to a non-pregnant female (D). (E) Comparison between the AgRP fluorescent signal intensity of non-pregnant female mice with *ad libitum* access to regular chow, pregnant mice with *ad libitum* access to regular chow, and pregnant mice with 24h hour food intake matched to a non-pregnant female (n=5-6) (F-G) Confocal images of POMC mRNA (green) expression and DAPI (blue) in Arc of female non-pregnant mouse with *ad libitum*access to food (F), pregnant female mouse with *ad libitum* access to food (G), and pregnant mouse with 24h hour food intake matched to a non-pregnant female (H). (I) Comparison between the POMC fluorescent signal intensity of non-pregnant female mice with *ad libitum* access to regular chow, pregnant mice with *ad libitum* access to regular chow, and pregnant mice with 24h hour food intake matched to a non-pregnant female (n=5-6). (J-K) Confocal images of AgRP (red) and fos (white) mRNA expression and DAPI (blue) in Arc of female non-pregnant mouse with *ad libitum* access to food (J) and non-pregnant mice fasted for 24h (K). (L) Comparison between the number of AgRP expressing cells colocalized with fos in non-pregnant female mice with *ad libitum* access to food or fasted for 24h (n=5-10). (M-N) Confocal images of AgRP (red) and fos (white) mRNA expression and DAPI (blue) in Arc of pregnant mouse with *ad libitum* access to food (M) and pregnant mouse with 24h hour food intake matched to a non-pregnant female (N). (O) Comparison between the number of AgRP expressing cells colocalized with fos in non-pregnant female mice with *ad libitum* access to food, pregnant mouse with ad libitum access to food, and pregnant mouse with 24h hour food intake matched to a non-pregnant female (n=5-11). (P-Q) Confocal images of POMC (red) and fos (white) mRNA expression and DAPI (blue) in Arc of female non-pregnant mouse with *ad libitum* access to food (P) and non-pregnant mouse fasted for 24h (Q). (R) Comparison between the number of POMC expressing cells colocalized with fos in non-pregnant female mice with *ad libitum* access to food or fasted for 24h (n=5-10). (S-T) Confocal images of POMC (red) and fos (white) mRNA expression and DAPI (blue) in Arc of pregnant mouse with *ad libitum* access to food (S) and pregnant mouse with 24h hour food intake matched to a non-pregnant female (T). (U) Comparison between the number of POMC expressing cells colocalized with fos in non-pregnant female mice with *ad libitum* access to food, pregnant mouse with *ad libitum* access to food, and pregnant mouse with 24h hour food intake matched to a non-pregnant female (n=5-11). (V) Post-pregnancy experiment timeline (W-Y) Confocal images of AgRP (red) and fos (green) mRNA expression and DAPI (blue) in Arc of female non-pregnant (W), pregnant (X), and 5 days after pregnancy mice (Y). (Z) Comparison between the number of AgRP expressing cells colocalized with fos in non-pregnant, pregnant and post-pregnancy mice (n=5-11). (AA-AC) Confocal images of POMC (red) and fos (green) mRNA expression and DAPI (blue) in Arc of female non-pregnant (AA), pregnant (AB), and 5 days after pregnancy mice (AC). (AD) Comparison between the number of POMC expressing cells colocalized with fos in non-pregnant, pregnant and post-pregnancy mice (n=5-11). Data are presented as mean values ± SEM. For panels E and I, statistical significance was tested by one-way ANOVA and Tukey’s multiple comparisons test. For panels L and R, statistical significance was tested by unpaired t test with Welch’s correction. For panels Z and AD, statistical significance was tested by 2way ANOVA and Tukey’s multiple comparisons test. For all panels *p\** < 0.05, *p\*\** < 0.01, *p\*\** < 0.001. Data points represent individual mice.

### Pregnancy increases AgRP neuron activity

Although AgRP mRNA levels were changed during pregnancy (**Fig. 2B-E**), this data does not provide information related to the neuronal activity levels of AgRP neurons during pregnancy. Therefore, we next utilized RNAscope in situ hybridization for the immediate early gene and marker of neuronal activity fos, together with AgRP or POMC mRNA detection to characterize the relative neuronal activity of AgRP and POMC neurons in non-pregnant and third trimester pregnant mice (**Fig. 2J-U**). As an additional control, in separate groups of mice, we measured fos activity in AgRP and POMC neurons following a 24-hour fast. Consistent with prior studies,^25, 26^ fasting increased the number of AgRP neurons co-expressing fos (**Fig. 2J-L**). Like fasting, pregnant mice also had a significantly greater number of AgRP neurons containing fos, relative to virgin *ad libitum* fed mice (**Fig. 2M-O**). Strikingly, matching the food intake of the pregnant mice to the levels observed in non-pregnant mice resulted in a similar recruitment of AgRP neuron activity as observed in 24-hour fasted virgin mice (**Fig. 2N-O**), indicating a substantial stimulatory effect of pregnancy on AgRP neuron activity.

In addition to AgRP neurons, an additional population of non-AgRP expressing neurons in the arcuate neurons contain vGAT, drastically increase food intake when activated, and are proposed to contribute to the anorexic actions of leptin.^27, 28^. Therefore, to determine if pregnancy specifically increases activity of AgRP neurons, or if pregnancy results in more widespread activation of many orexigenic cell types in the arcuate nucleus, we also characterized fos expression in arcuate vGAT neurons in pregnant and non-pregnant mice (**Ext Fig. 1**). However, in contrast to AgRP neurons, pregnancy did not significantly alter fos expression in arcuate vGAT neurons (**Ext Fig. 1A-C**). Therefore, pregnancy specificity increases the neuronal activity of arcuate AgRP neurons, without broadly increasing the neuronal activity in most orexigenic arcuate populations.

### Pregnancy decreases POMC neuron activity

Since we observed reduced POMC mRNA levels during pregnancy (**Fig. 2F-I**), we next quantified the expression of fos in POMC neurons in non-pregnant mice following a 24 hour fast and during the third trimester of pregnancy. As expected, fasting decreased the number of POMC neurons co-expressing fos mRNA, validating the effectiveness of this approach for detecting changes in POMC activity (**Fig. 2P-R**). Like fasting, pregnant mice exhibited a decrease in the number of POMC neurons co-expressing fos compared to non-pregnant mice (**Fig. 2S-U**). The inhibitory response of POMC neurons during pregnancy was indistinguishable from the inhibitory response of POMC neurons to a 24 hour fast, indicating a significant inhibitory effect of pregnancy on POMC neuron activity. Further, matching the food intake of pregnant mice to the levels observed in non-pregnant mice further reduced the number of POMC neurons containing fos (**Fig. 2S-U**).

We next tested if the inhibitory response of arcuate POMC neurons during pregnancy was specific to POMC neurons in the arcuate nucleus, or if other anorexia promoting neurons in the arcuate nucleus also exhibit reduced activity during pregnancy. To answer this question, we next quantified fos expression in arcuate neurons containing vGLUT2 (**Ext Fig. 1D-F**), since these cells only partially overlap with POMC neurons and have been reported to rapidly suppress feeding behavior.^29^ Both pregnant and non-pregnant mice exhibited similar levels of fos expression in arcuate vGLUT2 neurons (**Ext Fig. 1D-F**). Thus, pregnancy concordantly increases the neuronal activity of AgRP neurons while reducing the activity of POMC neurons, likely promoting increased feeding behavior.

### Pregnancy-induced changes in melanocortin circuitry partially reverse following parturition

Although AgRP neuron activity increases and POMC neuron activity decreases during pregnancy, it remains unclear if these changes are transient in nature or if they persist following pregnancy. Thus, we repeated RNAscope analysis for AgRP and POMC colocalization with fos in non-pregnant mice, during the third trimester of pregnancy, and 5 days following pregnancy (**Fig. 2V**). For these experiments, in the post-pregnancy group, pups were removed from the dam on the day of parturition to eliminate the potential confound of lactation on melanocortin circuit activity. As previously demonstrated, pregnancy significantly increased fos expression in AgRP neurons (**Fig. 2W-Z**). Interestingly, fos expression in AgRP neurons was completely reversed to non-pregnant levels following pregnancy, indicating that the increased AgRP neuron activity is transient and specific to the state of pregnancy (**Fig. 2Y-Z**). In contrast, although POMC neuron activity decreased during pregnancy, POMC activity did not completely return to baseline following pregnancy (**Fig. 2AA-AD**), suggesting differences in the temporal dynamics of AgRP and POMC neuron activity during the perinatal period.

### Pregnancy enhances the sensory response of AgRP neurons to palatable food

AgRP neurons are rapidly modulated by the sensory detection of food in hungry mice.^30–, 32^ Since *ex vivo* RNAscope analysis cannot capture these rapid changes in AgRP neuron activity in response to food presentation, we next performed *in vivo* fiber photometry from AgRP neurons in response to an acute fast in both non-pregnant and pregnant mice. AgRP-Cre mice were injected with a Cre recombinase dependent version of the genetically encoded calcium indicator GCAMP6s into the arcuate nucleus, and a fiber optic cannula was inserted into the arcuate nucleus for recording changes in fluorescence (**Fig. 3A**). First, we measured the dynamics of AgRP neuron activity to the presentation of a regular chow food pellet following a 45 minute fast during the dark period in all mice (**Fig. 3B**). After a 45-minute fast, mice were presented first with a non-accessible caged chow pellet for five minutes to assess the sensory response to the presentation of chow, followed by the removal of the cage covering the chow for five additional minutes to measure the AgRP neuronal response to accessible food (**Fig. 3B**). On the same day, food was again removed from the cage for an additional 5 hours and the response of AgRP neurons to both caged and accessible food was again measured in all mice (**Fig. 3B**). Following these initial calcium imaging experiments, all mice were either mated with a male mouse to induce pregnancy or paired with a female mice as a control, and the identical experiment was performed again during the third trimester of pregnancy (or the equivalent time-period in non-pregnant “time-matched” control mice). Following 45 minutes of food deprivation, AgRP neuron activity was equivalently reduced in the pre-pregnancy and pregnancy period in the seconds following the presentation of the “caged” chow, and in the seconds following access to the food pellet (**Fig. 3B**). AgRP neuron activity was also equivalently responsive to the presentation of a “caged” food pellet and upon access to the food following 6 hours of food deprivation during both the pre-pregnancy period and during pregnancy (**Fig. 3B**). Thus, although the basal activity of AgRP neurons is increased in pregnancy (**Fig. 2**), pregnancy does not alter the acute responsivity of AgRP neurons to the presentation of standard chow in food deprived mice (**Fig. 3C-F**).

**Figure 3:**
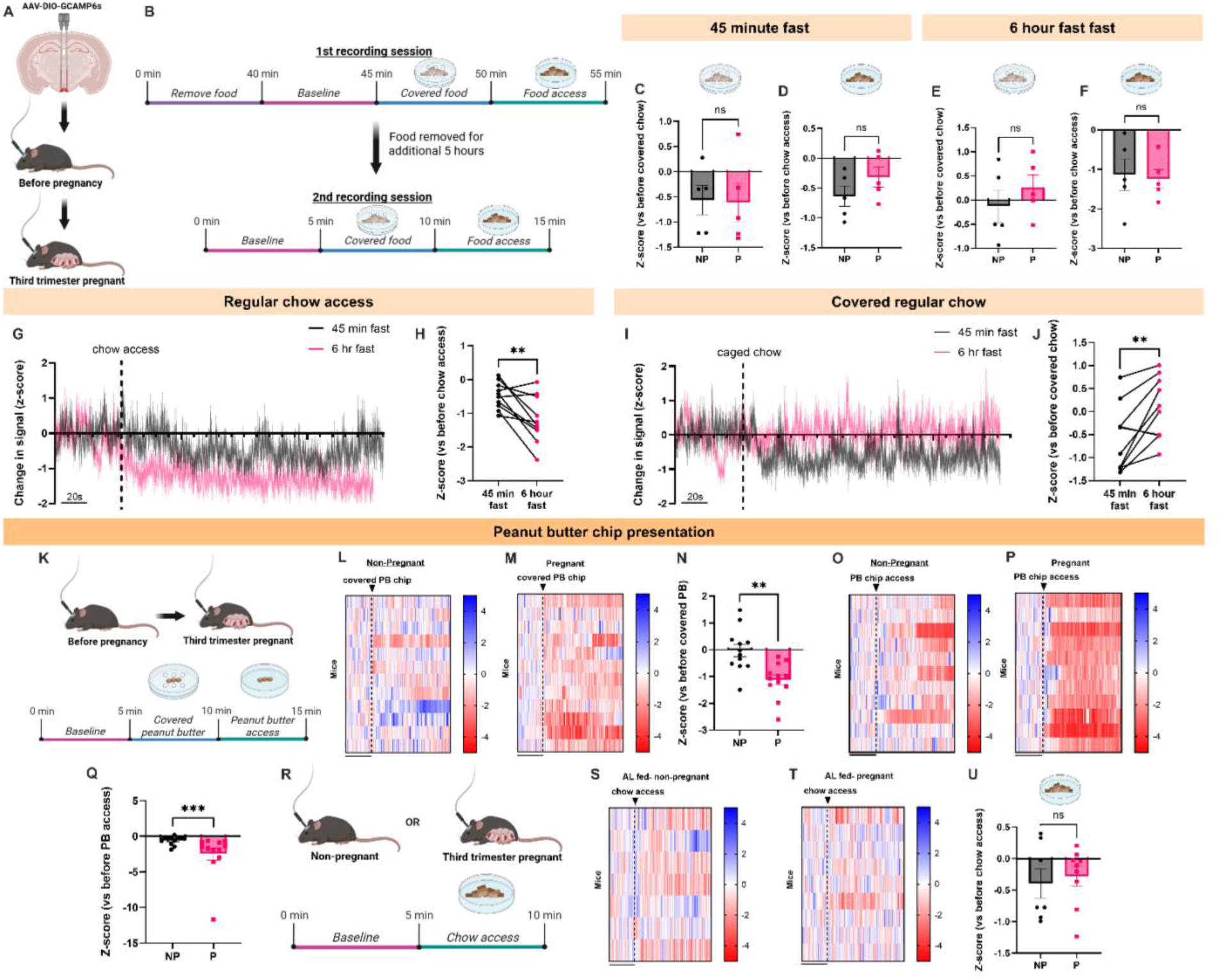
AgRP neurons have impaired response to palatable food during pregnancy. (A) Schematics of the AgRP-cre/GCAMP6a fiber photometry experiments. (B) Timeline of the recording sessions before (NP = non-pregnant) and during third trimester of pregnancy (P = pregnant). (C-D) Before and during pregnancy z-score comparison of the AgRP neuronal response after 45 minutes of fasting to the presence of covered regular chow (C) and with access to the food (D) (n=5 mice). (E-F) Before and during pregnancy z-score comparison of non-pregnant and pregnant mice AgRP neuron response, 6 hours after fasting, with the presence of covered regular chow (E) and with access to the food (F) (n=5 mice). (G) Change in the signal of the AgRP neurons during the covered regular chow phase before and during pregnancy, after 45 minutes or 6 hours of fasting (n=10 trials from 5 mice; 2 trials/mouse). (H) Comparison of the AgRP neurons calcium activity z-score of non-pregnant and pregnant mice with the presence to covered regular chow after fasting for 45 minutes or 6 hours (n=10 trials from 5 mice; 2 trials/mouse). (I) Change in the signal of the AgRP neurons during access to regular chow before and during pregnancy, after 45 minutes or 6 hours of fasting (n=10 trials from 5 mice; 2 trials/mouse). (J) Comparison of the AgRP neuronal calcium activity of non-pregnant and pregnant mice with access to regular chow after fasting for 45 minutes or 6 hours (n=10 trials from 5 mice; trials/mouse). (K) Timeline of the recording sessions with palatable food before and during third trimester of pregnancy. (L-M) Heat map of AgRP calcium activity recordings during presence of covered peanut butter chip (PB) for non-pregnant (L) and pregnant (M) mice (scale bars = 60s, n=12 mice). (N) Comparison of the AgRP neuronal calcium activity of non-pregnant and pregnant mice before and after the presence of the covered peanut butter chip (n=12 mice). (O-P) Heat map of AgRP calcium activity recordings during the access to peanut butter chip for non-pregnant (O) and pregnant (P) mice (scale bars = 60s, n=11 mice). (Q) Comparison of the AgRP neuronal calcium activity of non-pregnant and pregnant mice before and after access to peanut butter chip (n=12 mice). (R) Timeline of the recording sessions with regular chow before and during third trimester of pregnancy. (S-T) Heat map of AgRP calcium activity recordings during access to regular chow for non-pregnant (S) and pregnant (T) mice (scale bars = 60s, n=7-9). (U) Comparison of the AgRP neuronal calcium activity of non-pregnant and pregnant mice after access to regular chow (n=7-9). Data are presented as mean values ± SEM. For panels A-Q, statistical significance was tested by paired t-test. For panels R-U, statistical significance was tested by unpaired t-test. For all panels *p\** < 0.05, *p\*\** < 0.01, *p\*\** < 0.001. Data points represent individual mice.

Interestingly, we noticed in our prior experiments that while AgRP neuron activity was inhibited following the sensory detection of food after a 45-minute fast, AgRP neurons did not respond to the sensory detection of food following 6 hours of food deprivation (**Fig. 3G-H**). These results are surprising as 6 hours of food deprivation may be expected to increase AgRP neuron activity to a greater extent than 45 minutes of food deprivation. Therefore, we next reanalyzed the difference between the change in calcium signal to the presentation of caged food and accessible food in the same mice following a 45 minute fast and a 6 hour fast. As expected, a greater decrease in calcium signal was detected following access to chow after a 6-hour fast than a 45-minute fast, consisted with increased hunger (**Fig. G-H**). In contrast, while AgRP neurons were inhibited following the presentation of a caged food pellet after a 45-minute fast, these same neurons were no longer inhibited by the presentation of a caged food pellet on the same day following 6 hours of food deprivation (**Fig. 3I-J**). These results suggest that the sensory response of AgRP neurons to the detection of food is sensitive to single trial learning,^33^ such that AgRP neurons are no longer rapidly inhibited following caged food presentation once the mouse learns that this food will not be accessible.

Next, we tested the neuronal responsivity of AgRP neurons in non-pregnant and pregnant mice to the presentation of palatable peanut butter (PB) chips in the *ad libitum* fed state. Mice were first acclimated to the PB chip for two days prior to testing by providing one small PB chip (approximately 0.3g) in the mouse’s home cage for two days prior to fiber photometry testing. As previously described, on the fiber photometry test day, mice were sequentially presented with a caged PB chip, following by the removal of the cage and access to the PB chip (**Fig. 3K**). The same experiment was subsequently repeated during the third trimester of pregnancy or the equivalent time-period in non-pregnant mice. Presentation of a caged PB chip resulted in a significantly greater inhibitory response in AgRP neurons in pregnant mice than observed in the same mice prior to pregnancy (**Fig. 3L-N**). In contrast, no difference in the response to PB chip presentation was detected in non-pregnant control animals at both time-points (**Ext Fig. 2A**). Presentation of a covered peanut butter chip also trended towards producing a more significant inhibitory response in pregnant mice than non-pregnant mice on the same test day (**Ext Fig. 2B**). Following PB chip access, the magnitude of AgRP neuronal inhibition to an accessible PB chip was also enhanced in the pregnancy period compared to non-pregnant mice (**Fig. 3O-Q**), and in pregnant mice compared to non-pregnant mice on the same test day (**Ext Fig. 2D**). No difference in the responsivity of AgRP neurons to PB chip access was detected at both time-periods (i.e. pre-pregnancy and pregnancy period) in non-pregnant control mice (**Ext Fig. 2C**). The enhanced inhibitory response to food presentation in pregnant mice is specific to the presentation of palatable, calorically dense chow as presentation of a standard chow food pellet to sated mice did not alter AgRP neuron activity in both non-pregnant and pregnant mice (**Fig. 3R-U**). Therefore, pregnancy enhances the inhibitory response of AgRP neurons to the presentation of palatable food in the sated state.

### Inhibition of AgRP neurons reverses pregnancy-induced hyperphagia

Having established that pregnancy is associated with increased AgRP neuron activity (**Fig. 2M-O**), we next sought to determine if AgRP neurons are required for promoting hyperphagia during pregnancy. To test for a causal role for AgRP neurons in pregnancy-induced hyperphagia, we utilized chemogenetic inhibition approaches to selectively inhibit AgRP neurons in non-pregnant and pregnant mice. The chemogenetic inhibitor hM4Di was expressed in AgRP neurons via Cre-recombinase dependent AAV injections into the arcuate nucleus in AgRP-Cre mice (**Fig. 4A-C**). Following viral injections, we mated half of the experimental mice with male mice to induce pregnancy, while the remaining animals were paired with female mice as non-pregnant control animals. Consistent with recent reports indicating a negligible effect of acute inhibition of AgRP neurons in *ad libitum* fed mice,^28, 34, 35^ CNO-mediated inhibition of AgRP neurons (1mg/kg, i.p.) did not significantly reduce feeding in non-pregnant animals (**Fig. 4D**). In contrast, inhibition of AgRP neurons significantly reduced food intake in pregnant mice to levels which were indistinguishable from non-pregnant control mice (**Fig. 4D**). This reduction in feeding was mediated by reduced meal size (**Fig. 4E**), with no effect on meal number (**Fig. 4F**). In contrast to pregnant mice, inhibition of AgRP neurons did not alter meal size or meal number in non-pregnant animals (**Fig. 4E-F**). Thus, the increased feeding and meal size associated with pregnancy requires AgRP neuron activity.

**Figure 4:**
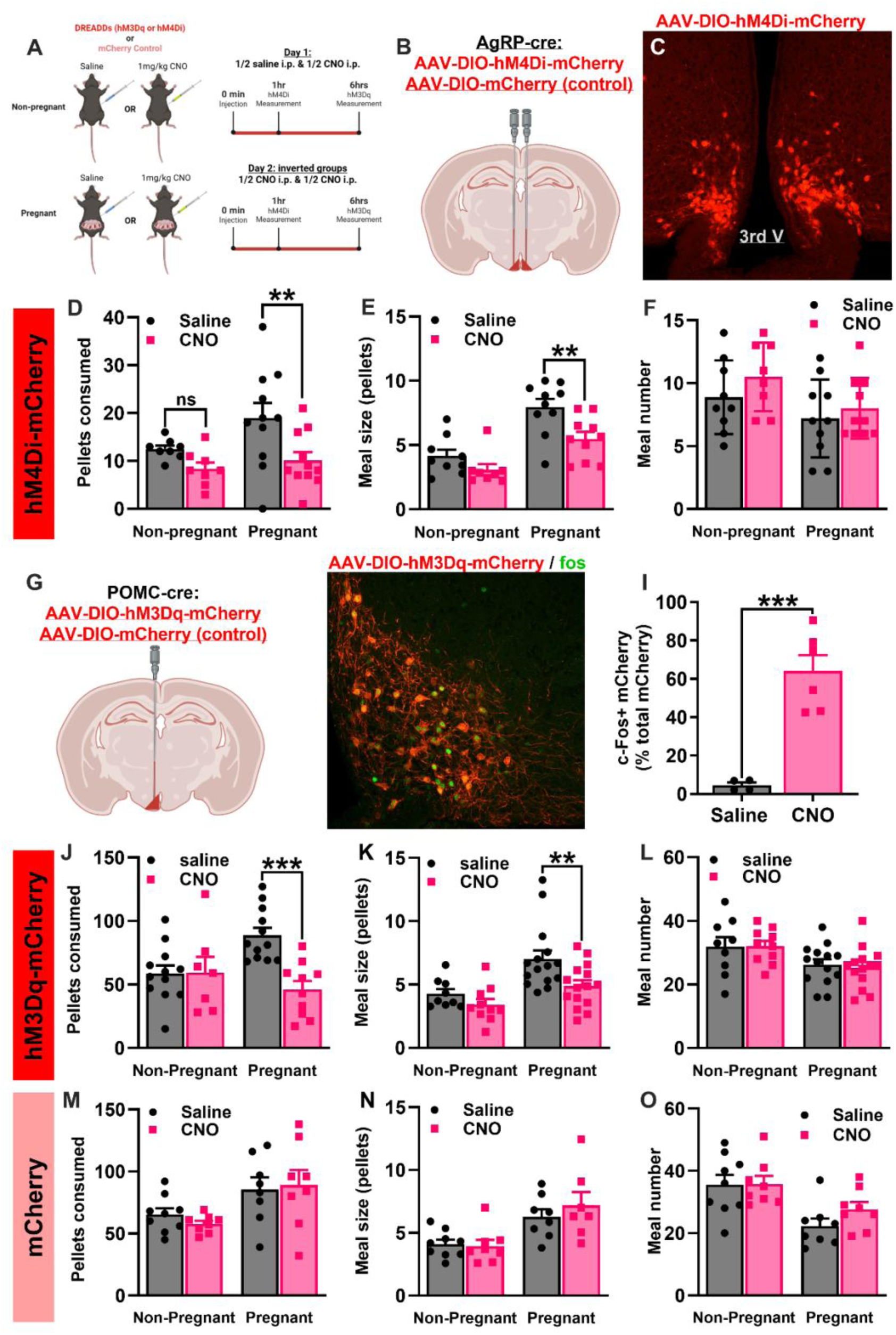
Inhibition of AgRP neurons or activation of POMC neurons reverses pregnancy hyperphagia. (A) Chemogenetic experiment schematics and timeline. (B) Schematics of the stereotaxic surgery and viruses utilized in the AgRP-cre/hM4Di experiment. (C) Example image of the viral location in Arc. (D-F) Comparison between non-pregnant and pregnant AgRP/hM4Di mice after saline or CNO intraperitoneal administration of pellets consumed in 1 hour (D), number of pellets consumed per meal (E), and number of meals following saline or CNO injections in non-pregnant and pregnant mice expressing hM4Di in AgRP neurons (n=8-11). (G). Schematics of the stereotaxic surgery and viruses utilized in the POMC-cre/hM3Dq experiment. (C) Example image of the viral location (red) and fos protein (green) in Arc (H). (I) Analysis of the percentage of POMC cells expressing fos after saline or CNO administration (n=4-6). (J-L) Comparison between non-pregnant and pregnant POMC/hM3Dq mice after saline or CNO intraperitoneal administration of pellets consumed in 6 hours (J), number of pellets consumed per meal (K), and number of meals in non-pregnant and pregnant mice expressing hM3Dq in POMC neurons following saline or CNO injections (L) (n=7-15). (M-O) Number of pellets consumed in non-pregnant and pregnant POMC/mCherrry mice after saline or CNO injections in 6 hours (M), number of pellets consumed per meal (N), and number of meals consumed (O) (n=8-9). Data are presented as mean values ± SEM. For panels D-F, J-L, and M-O, statistical significance was tested by 2way ANOVA and Šídák’s multiple comparisons test. For panel I, statistical significance was tested by Unpaired t test with Welch’s correction. For all panels *p\** < 0.05, *p\*\** < 0.01, *p\*\** < 0.001. Data points represent individual mice.

### Activation of POMC neurons reverses the hyperphagia of pregnancy

In contrast to AgRP neurons, arcuate nucleus neurons expressing pro-opiomelanocortin (POMC) are activated by energy sufficiency and release the melanocortin receptor agonist alpha-melanocyte stimulating hormone (aMSH) to suppress feeding.^10^ Our prior results (**Fig. 2**) indicate that both mRNA levels of POMC and fos activity in POMC neurons is lower in pregnant mice, suggesting that reduced POMC neuronal activity contributes to hyperphagia in pregnancy. Therefore, we hypothesized that pregnancy may reduce POMC neuron activity as a mechanism to promote increased feeding. If this is the case, we reasoned that pregnant mice may be more sensitive to stimulation of POMC neurons than non-pregnant animals, since the baseline activity of these neurons is low during pregnancy. To test this hypothesis, we expressed the chemogenetic activator hM3Dq in POMC neurons^36^ (**Fig. 4G**), allowing for selective activation of POMC neurons in non-pregnant and pregnant mice. Administration of the DREADD agonist CNO (1mg/kg, i.p.) increased the expression of the immediate early gene cfos in POMC neurons (**Fig. 4H-I**), indicating successful stimulation of POMC neurons following CNO injections. As observed in prior studies,^35, 37^ chemogenetic activation of arcuate POMC neurons did not acutely alter feeding behavior in non-pregnant *ad libitum* fed mice (**Fig. 4J**). In contrast, activation of POMC neurons drastically reduced feeding in pregnant mice by approximately 50 percent (**Fig. 4J**). Like inhibition of AgRP neurons (**Fig. 4E-F**), activation of POMC neurons selectively reduced meal size in pregnant mice (**Fig. 4K**), without altering the number of meals (**Fig. 4L**). Importantly, administration of CNO did not alter food intake or feeding structure in both non-pregnant and pregnant animals expressing control mCherry virus in POMC neurons (**Fig. 4M-O**), indicating that the enhanced sensitivity to POMC neuron stimulation was not mediated by off-target effects of CNO or viral expression. Together, chemogenetic studies indicate that pregnancy is associated with reduced basal melanocortin tone, resulting in an enhanced anorexic response to exogenous stimulation of the central melanocortin pathway (via either AgRP neuron inhibition or POMC neuron activation).

### Pregnant mice are hypersensitive to stimulation of central melanocortin circuits

Given that AgRP and POMC neurons can regulate feeding by inhibiting (via AgRP) or activating (via aMSH) downstream melanocortin receptors, we next tested if pregnancy is associated with an altered response to melanocortin receptor stimulation. We hypothesized that if pregnancy results in reduced activation of downstream MC4R (i.e. via elevated AgRP neuron activity and reduced POMC neuron activity), then pharmacological stimulation of melanocortin receptors may be more potent in pregnant than non-pregnant mice. To test this hypothesis, we quantified the feeding response of non-pregnant and pregnant mice to administration of the melanocortin receptor agonist setmelanotide (i.p.; 100ug/kg and 300ug/kg; **Fig. 5A**). Consistent with reduced melanocortin tone during pregnancy, we observed a significantly enhanced anorexic response to exogenous melanocortin receptor stimulation in pregnant mice (**Fig. 5B-C**). Importantly, this effect was not due to changes in CNS penetrance or pharmacokinetic changes associated with pregnancy as a similar phenotype was also observed following intra-cerebral ventricular administration of the same dosage of setmelanotide (vs aCSF) to pregnant and non-pregnant mice (**Fig. 5D**).

**Figure 5:**
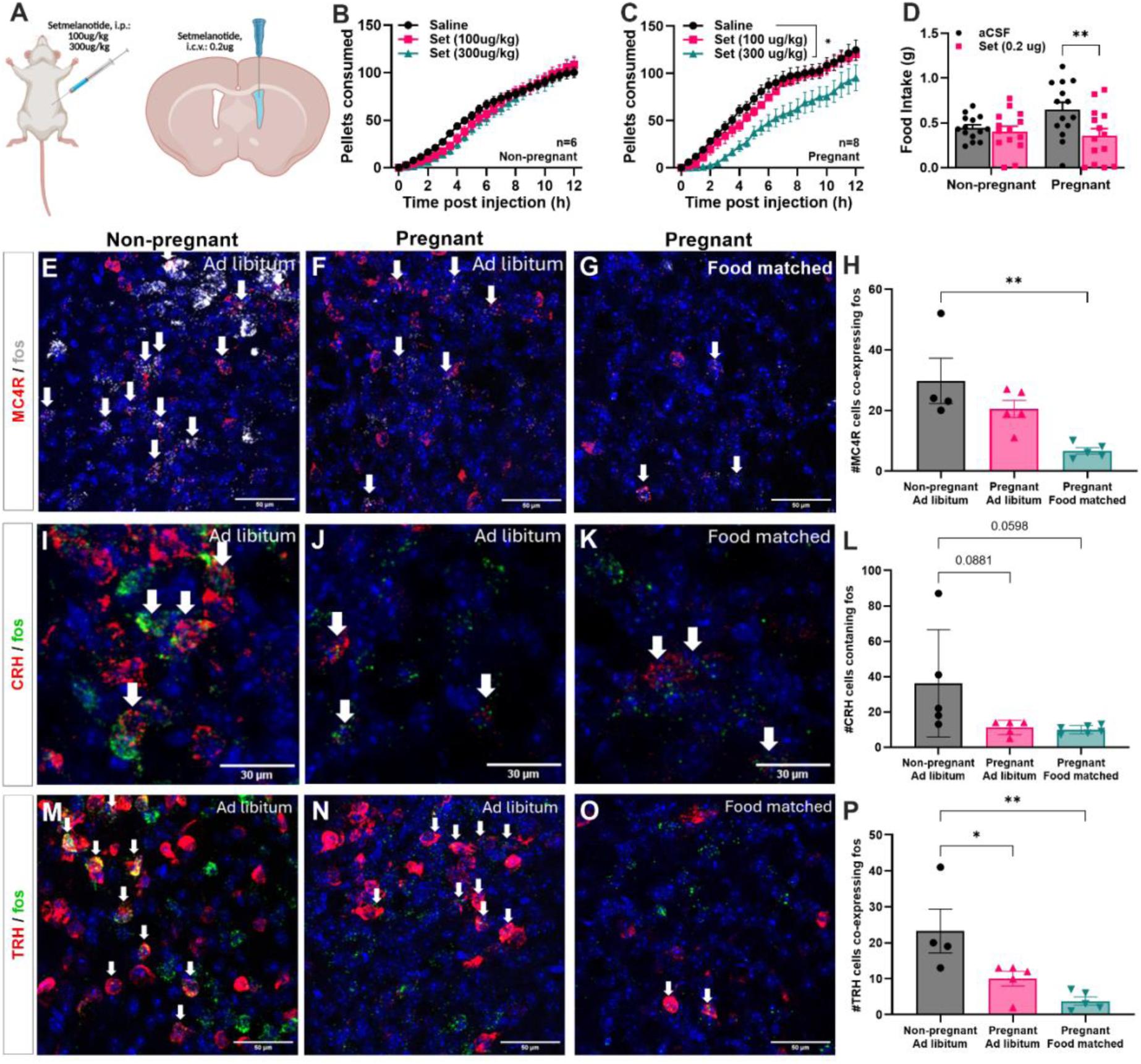
Pregnant mice are hypersensitive to stimulation of melanocortin receptors. (A) Schematics showing setmelanotide administration experiment via intraperitoneal (i.p) or intracerebroventricular (i.c.v) injections in non-pregnant or pregnant animals. (B-C) Comparison of pellets consumed after i.p administration of saline, 100ug/kg, or 300ug/kg setmelanotide in non-pregnant (B) or pregnant (C) female (D) mice (n=6-8). (D) Food intake, in grams, in non-pregnant and pregnant mice after i.c.v. administration of artificial cerebral spinal fluid (aCSF) or 0.2ug of setmelanotide (set.) (n=14) (E-G) Confocal images of MC4R (red) and fos (white) mRNA expression and DAPI (blue) in the PVN of pregnant mouse with *ad libitum* access to food (E) and pregnant mouse with *ad libitum* access to food (F), and pregnant animals with 24h hour food intake matched to a non-pregnant female (G). (H) Comparison between the number of MC4R expressing cells colocalized with fos in non-pregnant female mice with *ad libitum* access to food, pregnant mouse with *ad libitum* access to food, and pregnant mouse with 24h hour food intake matched to a non-pregnant female (n=4-5). (I-K) Confocal images of CRH (red) and fos (green) mRNA expression and DAPI (blue) in the PVN of pregnant mouse with *ad libitum* access to food (I) and pregnant mouse with *ad libitum* access to food (J), and pregnant animals with 24h hour food intake matched to a non-pregnant female (K). (L) Comparison between the number of CRH expressing cells colocalized with fos in non-pregnant female mice with *ad libitum* access to food, pregnant mouse with *ad libitum* access to food, and pregnant mouse with 24h hour food intake matched to a non-pregnant female (n=5-6). (M-O) Confocal images of TRH (red) and fos (green) mRNA expression and DAPI (blue) in the PVN of pregnant mouse with *ad libitum* access to food (M) and pregnant mouse with *ad libitum* access to food (N), and pregnant animals with 24h hour food intake matched to a non-pregnant female (O). (P) Comparison between the number of MC4R+ expressing cells colocalized with fos in non-pregnant female mice with ad libitum access to food, pregnant mouse with *ad libitum* access to food, and pregnant mouse with 24h hour food intake matched to a non-pregnant female (n=4-5). Data are presented as mean values ± SEM. For panels B and C, statistical significance was tested by one-way ANOVA and Dunnett’s multiple comparisons test. For panel D, statistical significance was tested by 2-way ANOVA and Šídák’s multiple comparisons test. For panels H, L, and P, statistical significance was tested by one-way ANOVA and Tukey’s multiple comparisons test. For all panels *p\** < 0.05, *p\*\** < 0.01, *p\*\** < 0.001. Data points represent individual mice.

We next tested if the enhanced response to anorexic drug treatment observed following setmelanotide administration was specific to melanocortin receptor stimulation or if pregnancy is associated with a more general state of increased responsivity to anorexic stimuli. In contrast to melanocortin receptor stimulation (**Fig. 5A-D**), pregnant mice exhibited a similar anorexic response as non-pregnant mice to the anorexic peptide’s peptide YY (PYY; **Ext Fig. 3A**), glucagon-like 1 receptor (GLP1R; **Ext Fig. 3B**) (liraglutide administration), and growth differentiation factor 15 (GDF15; **Ext Fig. 3C**). Thus, pregnancy enhances the anorexic response to melanocortin receptor stimulation, without an obvious change in the anorexic response to stimuli that do not directly target central melanocortin receptors.

To further confirm reduced melanocortin tone in pregnancy, we performed additional RNAscope experiments to characterize fos expression in downstream paraventricular hypothalamic (PVN) MC4R neurons in non-pregnant and pregnant mice (**Fig. 5E-G**). Like POMC neurons (**Fig. 2**), we observed a trend towards reduced fos expression in PVN MC4R neurons in *ad libitum* fed pregnant mice vs non-pregnant control animals (**Fig. 5H**). Compared to non-pregnant mice, fos expression in PVN MC4R neurons was drastically reduced in pregnant mice that were matched to the food intake levels of pregnant animals (**Fig. 5H**), indicating the pregnancy reduces the activity of PVN MC4R neurons when adjusting for caloric intake. To test if this reduced activity was specific to PVN MC4R neurons, or if other neuroendocrine cell types in PVN also reduce their activity during pregnancy, we performed similar analysis of PVN thyroid-releasing hormone (TRH), and corticotrophin-releasing hormone (CRH) neurons. Consistent with prior studies which have shown an attenuated activity of hypothalamic-pituitary adrenal axis (HPA) during pregnancy,^38, 39^ we observed a trend towards reduced activity of PVN CRH neurons during pregnancy in both *ad libitum* fed pregnant mice and in pregnant mice matched to the food intake levels of non-pregnant animals (**Fig. 5I-L**). Furthermore, fos expression in PVN TRH neurons was also reduced in pregnant mice compared to non-pregnant animals (**Fig. 5M-P**). This reduction in fos expression was further enhanced when pregnant mice were matched to the food intake levels of non-pregnant animals (**Fig. 5P**). Thus, pregnancy reduces neuronal activity in multiple behavioral (PVN MC4R) and neuroendocrine (PVN CRH and PVN TRH) cell types in the PVN, consistent with elevated food intake.

### Pregnancy alters the transcriptome of AgRP and POMC neurons towards the promotion of positive energy balance

To identify putative molecular mechanisms mediating altered AgRP and POMC neuron activity during pregnancy we next performed single cell resolution spatial transcriptomics (Xenium spatial transcriptomics) of 5000 protein coding genes in female non-pregnant mice fed *ad libitum*, fasted for 24 hours, or in the third trimester of pregnancy (**Fig. 6A and Ext Fig. 4**). To specifically characterize transcriptional changes within the arcuate nucleus we utilized neuroanatomical markers (i.e. AgRP and POMC) to virtually dissect and compare transcriptional changes within the arcuate nucleus across the three experimental conditions (**Fig. 6B)**. After clustering all cell types from the arcuate nucleus (**Fig. 6C-F**), we performed differential gene expression analysis for all cell types between *ad libitum* fed and fasted, and between *ad libitum fed* and the pregnant state. Neurons were the cell type with the highest number of differentially expressed genes (DEG) in response to both fasting and pregnancy, followed by astrocytes and tanycytes (**Fig. 6G and H**).

**Figure 6:**
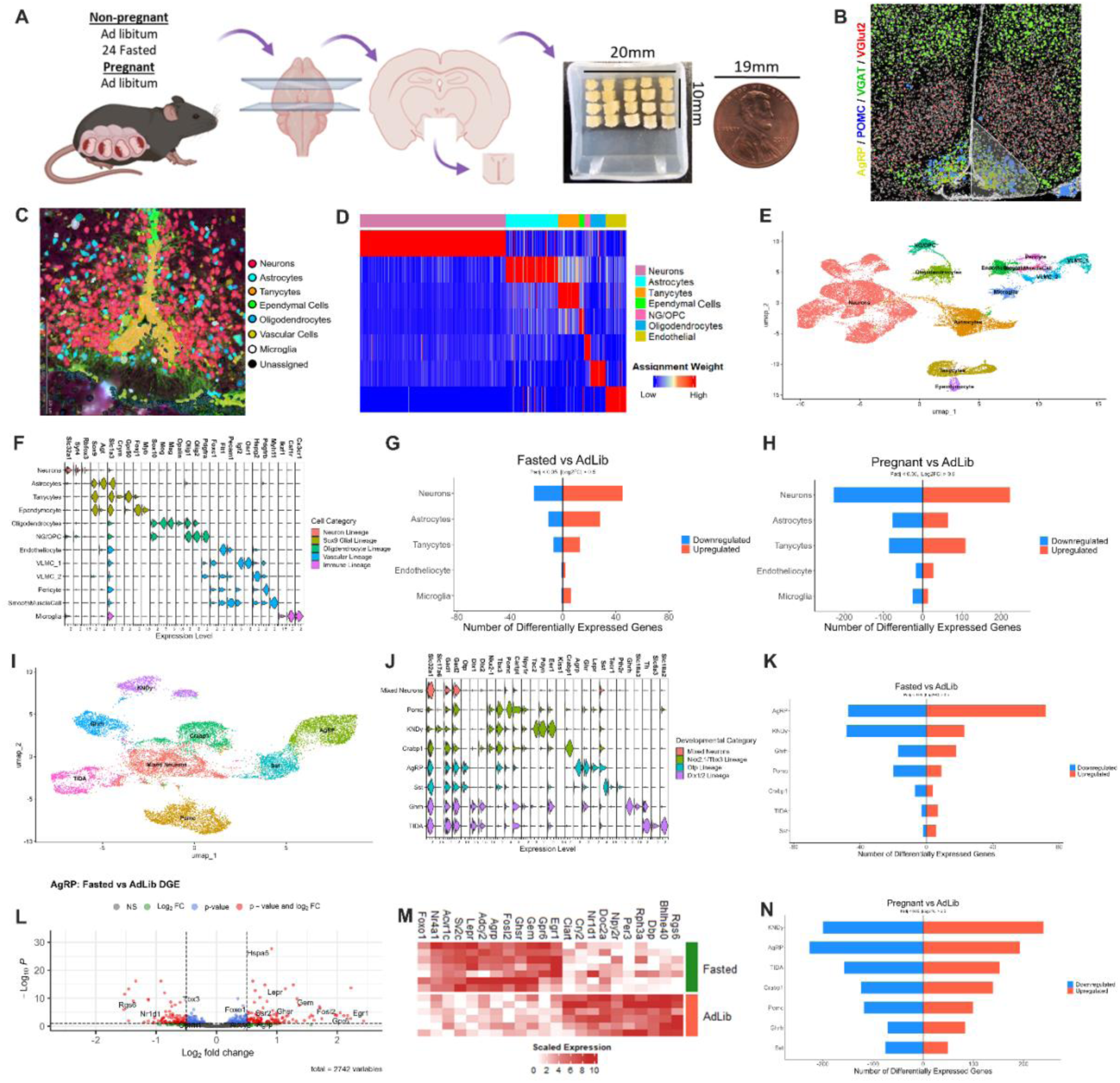
Transcriptomics changes in the arcuate nucleus are different in pregnancy and fasting conditions. (A) Schematic of sectioning and alignment of mediobasal hypothalami into a tissue array for spatial transcriptomic processing via *Xenium-in-situ* platform. (B) Representative coronal subsection of tissue following Xenium imaging. DAPI staining is presented in white, along with four mRNA transcripts with subcellular localization; AgRP (yellow), POMC (blue), VGAT (green), VGlut2 (red), each in canonical neuroanatomical regions *e.g.* VGAT in arcuate nucleus (ARC) and dorsomedial hypothalamus (DMH), and VGlut2 in ventromedial hypothalamus. Selection of the right division of the ARC is shown. (C) Example image of assigned neurons (red), astrocytes (cyan), tanycytes (orange), ependymal (green), oligodendrocytes (blue), vascular (yellow), microglia (white), and unassigned (black) cell types and their localization in an arcuate nucleus section (D) Heatmap of the assignment weight of the seven cell type clusters as determined via SingleR. (E) UMAP of SCTransformed transcriptomic data obtained for all ARC cells passing quality control. Labels are shown depicting manually curated cell-types. VLMC: Vascular leptomeningeal cells. (F) Violin plots of marker gene expression for each cell type in (E), cell types are organized into five lineages, based on co-expression of developmental transcription factors e.g. Sox9 in astrocytes, tanycytes, and ependymocytes. (G) Total number of genes differentially regulated between fasting and *ad libitum* fed conditions for the most abundant 5 cell-types. (H) Total number of genes differentially regulated between *ad libitum* pregnant and *ad libitum* fed conditions for the most abundant 5 cell-types. (I) UMAP of separately SCTransformed transcriptomic data obtained from ARC neurons across the three conditions. Labels are shown depicting manually curated neuronal cell types of the ARC. (J) Violin plot of marker gene expression for neurons in (I). Neurons are organized by developmental lineage via expression of transcription factors. (K) Total number of genes differentially regulated between fasting and *ad libitum* fed conditions for each neuron class (excluding mixed neurons). (L) Volcano plot of differential gene expression for AgRP neurons between fasting and *ad libitum* fed conditions. Known immediate early gene (IEG) and select genes with canonical roles in feeding are highlighted. (M) Heatmap of scaled, size-adjusted counts of some significantly differentially regulated genes for each sample (row) between fasting and *ad libitum* fed conditions. (N) Total Number of genes differentially regulated between pregnant and *ad libitum* fed conditions. Significance (G, H, K, L, N) was considered to be any gene with an absolute log-fold change greater than 0.5, and an adjusted P-value of less than 0.05. P-values were corrected using Benjamini-Hochberg protocol, and log-fold changes were scaled using apeglm shrinkage procedures. For all panels n=4-5 mice per group.

Since neurons contained the most differentially expressed genes in response to pregnancy, we utilized a panel of neuron marker genes to specifically re-cluster all the neurons within the arcuate nucleus across all experimental conditions (**Fig. 6I and J, Ext Data Fig. 4**). We identified most of the major neuron cell types in the arcuate nucleus, including arcuate AgRP, POMC, kisspeptin/dynorphic/neurokinin B (KNDy), tuberoinfundibular dopamine neurons (TIDA), CRABP1, growth hormone releasing hormone (GHRH), and somatostatin (SST) neurons (**Fig. 6I and J**). To determine the arcuate cell types most altered by fasting and pregnancy we compared the number of differentially expressed genes in response to fasting and pregnancy across all arcuate cell types. As expected, the hypothalamic AgRP neurons were the neuron cell type most transcriptionally altered by fasting (**Fig. 6K**). Consistent with prior reports,^40^ AgRP neurons exhibited approximately 5-10 fold more differentially expressed genes than POMC neurons in response to fasting, while other arcuate cell types were much less altered by fasting (**Fig. 6K**). Importantly, we observed upregulation of many genes previously validated to be upregulated by fasting in AgRP neurons (i.e. LepR, GHSR, GPR6, Foxo1, and multiple immediate early genes indicating increased neuron activity), validating the effectiveness of this approach for detecting transcriptional changes in specific arcuate cell types (**Fig. 6L and M**).

Next, we compared the number of differentially altered genes in each of the arcuate cell types in response to pregnancy (third trimester of pregnancy). Interestingly, AgRP neurons were also among the most transcriptionally altered by pregnancy, along with arcuate KNDy neurons and arcuate TIDA neurons, consistent with the important role of KNDy and TIDA neurons in controlling neuroendocrine and reproductive function (**Fig. 6N**). Both CRABP1 neurons and POMC neurons were also significantly altered during pregnancy, indicating that these neurons likely also contribute to changes in energy homeostasis during pregnancy (**Fig. 6N**). In general, pregnancy resulted in much more differentially expressed genes than fasting, perhaps reflecting the sustained and robust hormonal and neuroendocrine changes associated with this state (**Fig. 6K and N**).^1^ For example, we observed 5-10 fold more differentially expressed genes in AgRP and POMC neurons during pregnancy compared to fasting (**Fig. 6K and N**), indicating the drastic effect of pregnancy on the physiology of these neurons.

To further characterize the effects of fasting and pregnancy on the transcriptional state of AgRP neurons we performed principal component analysis and correlation matrixes on the arcuate AgRP neurons from the 25 independent arcuate sections (collected from 25 different mice) across the three experimental groups (*ad libitum* fed, fasted, and pregnancy; **Fig. 7A-D**). Principle component analysis (PCA) revealed that the transcriptional changes in AgRP neurons associated with these three states (*ad libitum* fed, fasted, pregnant) were clearly separated in principle component space, indicating that fasting and pregnancy produce distinct effects on the transcriptome of AgRP neurons. These findings were confirmed with correlation matrix results which demonstrated that the transcriptional effects of pregnancy on AgRP neurons were clearly distinct from both fasting and the *ad libitum* fed condition (**Fig. 7D**). In contrast, the AgRP transcriptome from different pregnant mice were highly similar from section to section, regardless of the specific mouse analyzed or the location (anterior or posterior) of the arcuate nucleus (**Fig. 7D**). To further elucidate putative molecular changes that may mediate the increased AgRP neuron activity during pregnancy we performed differential gene expression analysis between AgRP neurons from the *ad libitum* fed state and pregnancy. Among the genes most differently effected in AgRP neurons during pregnancy were genes associated with increased neuronal activity (Fosl2, Rheb), synaptic plasticity (Pak1, Pak3), and synaptic transmission (Syp, Acvr1c, Sv2c, Sv2a, Syn1, Gad2, Slc32a1, Foxo1) suggesting increased neuronal activity (**Fig. 7E and F**). To further confirm these findings, we performed post-hoc manual quantification of Gad2, Slc32a1, and Foxo1 in AgRP neurons from non-pregnant and pregnant mice, now analyzing the mRNA transcription levels per individual AgRP expressing cell. Consistent with transcriptomic analysis, both Gad2 and Slc32a1 levels were increased in AgRP neurons in pregnant mice, suggesting increased GABAergic transmission from AgRP neurons during pregnancy (**Fig. 7G-I**). Further, we observed increased expression of Foxo1 in AgRP neurons during pregnancy (**Fig. 7J**), which is also a phenotype observed in non-pregnant fasted females (**Fig. 7K**). Given that Foxo1 is a transcription factor which increases AgRP expression,^41–44^ these results are consistent with the increased AgRP expression observed during pregnancy in mice (**Fig. 2**).

**Figure 7:**
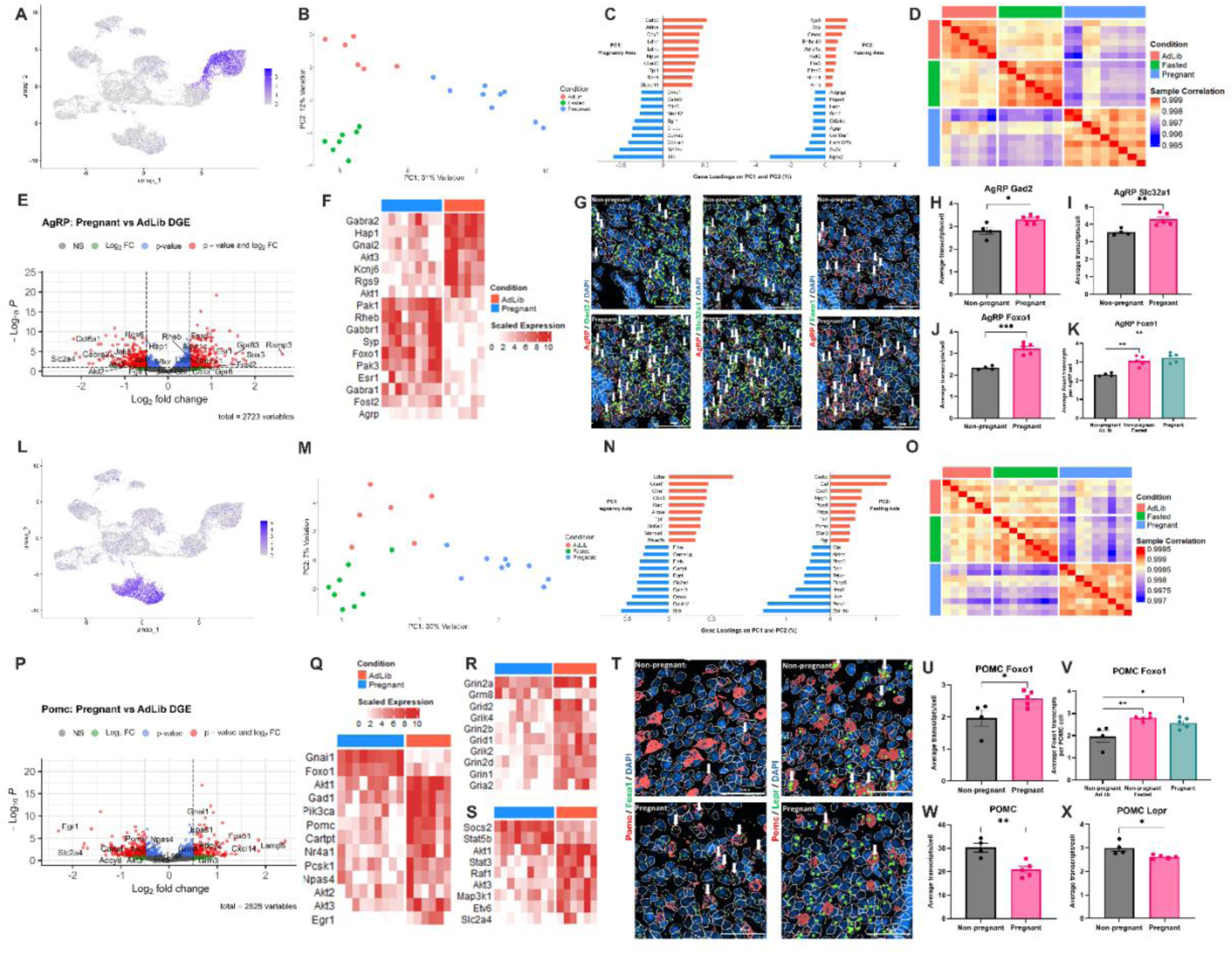
Pregnancy impairs both AgRP and POMC transcriptomics profile, differently from fasting, when compared to non-pregnant *ad libitum* females. (A) UMAP plot of arcuate neurons (as in Fig. 6F), colored by Pearson residuals of AgRP expression. (B) Principal components (PC) 1 and 2 of the AgRP joint-condition DESeq2 model, with *ad libitum* non-pregnant (red), fasted (green), and pregnant (blue) samples represented as individual dots. Percentage variation in the DESeq2 expression model explained by principal components is included along the axes. (C) Gene loadings for the top principal components from AgRP joint-condition DESeq2 model (B): PC1, which separates pregnant from non-pregnant conditions, and PC2, which broadly distinguishes *ad libitum* non-pregnant from fasted animals. The top 10 genes with the highest positive and negative loadings on each component are shown, with loadings expressed as a percentage of the total loading vector magnitude. (D) Sample-to-sample Pearson correlations of the AgRP joint-condition DESeq2 model, colored by condition: ad libitum (red), fasted (green), and pregnant (blue). (E) Volcano plot of differential gene expression for AgRP neurons between Pregnant and *ad libitum* conditions. (F) Heatmap of scaled size-adjusted counts of differentially regulated genes in AgRP neurons for each sample (row) between *ad libitum* (red) and Pregnant (blue) conditions. (G) Example images of AgRP (red) cells and their co-expression with Gad2, Slc32a1 or Foxo1 (green) with DAPI (blue) as nuclei marker. Arrows represent co-expressing nuclei of the AgRP and gene of interest. (H-K) Comparison between *ad libitum* non-pregnant and pregnant average number of transcripts counts in AgRP cells colocalized with Gad2 (H), Slc32a1 (VGAT) (I), Foxo1 (J), and Foxo1 comparing the 24-hour fasted non-pregnant females with ad libitum fed non-pregant females (K). (L) UMAP plot of arcuate neurons (as in Fig. 6F), colored by Pearson residuals of POMC expression. (M) PC1 and PC2 of the POMC joint-condition DESeq2 model, with *ad libitum* non-pregnant (red), fasted (green), and pregnant (blue) samples represented as individual dots. Percentage variation in the DESeq2 expression model explained by principal components is included along the axes. (N) Gene loadings for the top principal components from POMC joint-condition DESeq2 model, as in (B). (O) Sample-to-sample Pearson correlations of the POMC joint-condition DESeq2 model, colored by condition: ad libitum (red), fasted (green), and pregnant (blue). (P) Volcano plot of differential gene expression for POMC neurons between Pregnant and *ad libitum* non-pregnant conditions. (Q-S) Heatmaps of scaled size-adjusted counts of differentially regulated genes in POMC neurons for each sample (row) between *ad libitum* (red) and Pregnant (blue) conditions. (T) Example images of POMC (red) cells and their co-expression with Foxo1 or Lepr (green) with DAPI (blue) as nuclei marker. Arrows represent co-expressing nuclei of the POMC and gene of interest. (U-V) Comparison between *ad libitum* non-pregnant and pregnant mice of the average number of transcripts count in POMC cells which colocalize with Foxo1 (U), or Foxo1 expression in POMC neurons in 24-hour fasted non-pregnant females vs *ad libitum* fed non-pregnant females (V). (W and X) number of POMC transcripts in POMC neurons (W), and Lepr transcripts in POMC neurons in non-pregnant vs pregnant mice (X). In figures E and P, significance was considered to be any gene with an absolute log-fold change greater than 0.5, and an adjusted P-value of less than 0.05. P-values were corrected using Benjamini-Hochberg protocol, and log-fold changes were scaled using apeglm shrinkage procedures. For panels H-J, U, W, and X statistical significance was tested by unpaired t-test and for panels K and V was tested by 2-way ANOVA and Tukey’s multiple comparisons test, with *p\** < 0.05, *p\*\** < 0.01, *p\*\** < 0.001. Data points represent individual mice. For all panels n=4-5 per group.

We next further analyzed the transcriptional changes in POMC neurons during pregnancy by performing principal component analysis and correlation matrixes on the arcuate POMC neurons from the 25 arcuate sections across the three experimental groups (**Fig. 7L**). Consistent with the reduced gene expression changes associated with fasting in POMC neurons vs AgRP neurons, POMC neurons from fasted and *ad libitum* fed mice did not separate distinctly in principle component space (**Fig. 7M**). In contrast, POMC neurons from pregnant mice were clearly distinct in PCA space from POMC neurons in both the *ad libitum* fed state and the fasted state (**Fig. 7M and N**). Consistent with principal component results, we observed similar transcriptional changes in POMC neurons from all arcuate sections obtained from pregnant mice, while the transcriptional changes in POMC neurons from pregnant mice were significantly different than those of POMC neurons from both fasted and *ad libitum* fed conditions (**Fig. 7O**). We next utilized differential gene expression analysis from the POMC neurons from the *ad libitum* fed and pregnant state to identify putative molecular mechanisms mediating reduced POMC neuron activity during pregnancy. We observed many altered genes in POMC neurons during pregnancy suggestive of reduced neuron activity and synaptic transmission, including reduced expression of multiple immediate early genes (EgR1, Npas4), and genes involved in synaptic transmission/vesicle release (i.e. Nr4a1, Pcsk1, Carpt, POMC, Gad1) (**Fig. 7P and Q**). Interestingly, we also observed downregulation of many genes associated with postsynaptic glutamatergic transmission in POMC neurons during pregnancy (Gria2, Grid1, Grik4, Grin2b, Grin1, Grik2, Grin2d, Grid2, Grin2a), suggesting that reduced responsivity to glutamatergic inputs may contribute to the reduced POMC neuron activity observed during pregnancy (**Fig. 7R**).

Prior findings in rats suggest that pregnancy is associated with an impaired molecular response to downstream leptin and insulin signaling.^23, 45–48^ Since impaired leptin and insulin signaling in POMC neurons could provide one mechanism mediating reduced POMC activity during pregnancy, we also characterized the expression of downstream genes involved in leptin and insulin signaling in POMC neurons (i.e. JAK2/STAT3 and MAPK/PI3K related genes) from pregnant and non-pregnant mice (**Fig. 7S**). Despite high levels of insulin and leptin during pregnancy, ^23, 45–49^ we observed reduced expression of many genes associated with downstream leptin and insulin signaling in POMC neurons from pregnant mice (LepR, Stat3, Akt1, POMC, Map3k1, Akt3, Pik3ca, Etv6), suggesting that reduced functional leptin and insulin signaling may contribute to reduced activity of these neurons during pregnancy (**Fig. 7Q and S**). Further, as observed for AgRP neurons (**Fig. 7F and J**), POMC neurons from pregnant mice had increased expression per cell of the transcription factor Foxo1 (**Fig. 7T and U**), which was also observed in fasted non-pregnant females (**Fig. 7V**). Given that Foxo1 acts to reduce the expression of POMC by inhibiting downstream STAT3 signaling,^43, 44, 50^ these molecular results are consistent with reduced POMC expression in POMC neurons during pregnancy (**Fig. 2**). Finally, we confirmed the results of transcriptomic studies by manually quantifying the expression of POMC, Lepr, and Foxo1 in POMC neurons from pregnant and non-pregnant mice. Consistent with transcriptomic results, we observed reduced expression of both Lepr and POMC in POMC neurons, and increased expression of Foxo1 in POMC neurons during pregnancy (**Fig. 7T, W, and X**). Thus, pregnancy alters the transcriptome of POMC neurons in a manner which reduces POMC expression and promotes feeding.

## Discussion

The pregnancy period is associated with substantial increases in feeding behavior in mammals.^49^ This increased feeding is necessary to promote fetal growth and development and to facilitate energy storage (in the form of fat) for the subsequent energy demands of the lactational period.^49^ Despite this, the core neural mechanisms mediating an appropriate increase in feeding during pregnancy are incompletely understood. Here, consistent with prior reports,^51, 52^ we determine that increased feeding during pregnancy is driven by elevated meal size in mice, with pregnant mice drastically shifting their meal structure towards larger and more infrequent meals (**Fig. 1E-F**). Such a change may provide a mechanism to increase total calorie intake while limiting the amount of time dedicated towards foraging and food seeking.

Consistent with increased feeding, we identify both increased AgRP peptide expression and increased fos expression in AgRP neurons during pregnancy in mice (**Fig. 2B-E and 2M-O**). Conversely, POMC mRNA expression and fos activity in POMC neurons are reduced during pregnancy (**Fig. 2F-I and S-U**). These results are consistent with prior reports suggesting increased AgRP and reduced POMC mRNA expression during pregnancy in rodents.^21–23, 53^ However, other studies have reported no significant differences in AgRP and POMC expression during pregnancy in rodents.^22, 23^ Although the specific reasons for these inconsistencies are unknown, they may be due to species specific differences (i.e. rats vs mice), the specific pregnancy period recorded (i.e. second vs third trimester, etc), or technical differences associated with various approaches for quantifying mRNA expression.

Although increased feeding during pregnancy is adaptive for promoting fetal growth and development, in both humans and rodents’, pregnancy is associated with an increased risk for obesity and metabolic disorders later in life.^49^ Data presented here suggests that while AgRP activity is reversible following pregnancy, fos activity in POMC neurons remains slightly reduced following the pregnancy period (**Fig. 2W-AD**). This change may represent one molecular mechanism mediating increased obesity risk following pregnancy, although further work is required to precisely map long term changes in melanocortin circuit activity following pregnancy. However, given the critical role for AgRP and POMC neurons in energy homeostasis and glucose homeostasis,^10^ and the drastic changes in these neurons associated with pregnancy, future work is warranted to establish the causal role for these neurons in pregnancy associated disturbances in energy homeostasis, such as gestational diabetes and the increased risk for obesity and type 2 diabetes associated with childbirth.

Recent findings indicate that AgRP neurons are rapidly inhibited by the sensory detection of food, prior to food consumption.^31–33, 54^ Ultimately, food consumption durably inhibits these neurons to signal satiety and reduce food seeking behaviors.^33, 54^ Here, we demonstrate that pregnant and non-pregnant mice exhibit an equivalent inhibitory response to the presentation of standard chow following an acute fast (**Fig. 3C-F**). However, AgRP neurons from pregnant mice show an enhanced inhibitory response to the presentation of palatable chow (peanut butter chips) compared to non-pregnant animals (**Fig. 3L-Q**). Interestingly, this enhanced response is specific to the presentation of palatable food as AgRP neurons from both non-pregnant and pregnant mice do not respond to the presentation of standard chow in the *ad libitum* fed state (**Fig. 3S-U**). This data indicates that AgRP neurons are particularly sensitive to the presence of palatable, calorically dense foods during pregnancy. Given that both pregnant humans and mice have increased food cravings for palatable calorically dense food,^49, 55^ these data suggest that increased sensitivity of AgRP neurons to palatable foods may provide one cellular mechanism mediating increased palatable food cravings in pregnancy. Consistent with this hypothesis, AgRP neurons are more active in pregnant mice (**Fig. 2M-O**) and activation of AgRP neurons increases the mesolimbic dopamine response to palatable food consumption.^56, 57^ Furthermore, mesolimbic dopamine activity is enhanced in pregnant mice to promote increased high fat diet intake.^55^ Further work is ultimately required to determine the specific implications of the enhanced sensitivity of AgRP neurons to palatable food in pregnancy and the relationship this may have to increased palatable food cravings during pregnancy.

It is important to note that although prior reports suggest altered melanocortin circuit activity during pregnancy, ^21–23, 53^ the causal role for this system in increasing feeding during pregnancy was previously unknown. For example, changes in AgRP and POMC mRNA may occur secondarily to the altered neuroendocrine and metabolic changes associated with pregnancy and may not be causally implicated in promoting increased feeding during this period. However, chemogenetic results presented here indicate that both elevated AgRP neuron activity and reduced POMC neuron activity are required for promoting increased feeding during pregnancy (**Fig. 4**). Furthermore, pregnancy is associated with a drastic increase in the sensitivity of mice to stimulation of downstream melanocortin receptors (**Fig. 5**). These findings are reminiscent of syndromic obesity disorders characterized by hyperphagia resulting from reduced melanocortin tone.^58–60^ For example, although the MC4R agonist setmelanotide has limited efficacy in reducing feeding and body weight in humans with dietary obesity,^58^ setmelanotide is much more potent in patients with reduced melanocortin tone, reversing hyperphagia and obesity in patients with congenital leptin receptor or POMC mutations.^59, 60^ In contrast with results reported here, a prior report suggested that pregnant rats have reduced sensitivity to central administration of aMSH.^61^ Although the reasons for these differences are unknown, they may be due to species differences between rats and mice, differences between the pharmacology, kinetics, and sensitivity of the chemogenetic and pharmacology approaches used in this study vs this prior report, or the underlying energy state of the animals (i.e. *ad libitum* fed vs fasted). However, the pharmacology, chemogenetic, transcriptomic, and neural activity experiments presented here all support an important role for reduced melanocortin tone in promoting increased feeding during pregnancy in mice. Further work is required to determine the long-term effects of AgRP and POMC neuron activity on both normal and maladaptive feeding behavior during pregnancy, and to determine if increased AgRP neuron activity and reduced POMC neuron activity are required for a healthy and successful pregnancy.

Although the specific molecular mechanism(s) mediating altered AgRP and POMC neuron activity during pregnancy are unknown, spatial transcriptomic data presented here indicates that pregnancy drastically alters the transcriptional state of both AgRP and POMC neurons (**Fig. 6 and 7**). Pregnancy increases levels of the anorexic hormone’s leptin and insulin, which collectively increase POMC neuron activity and reduce AgRP neuron activity.^10, 26, 62–66^ Despite elevated levels of leptin and insulin, central responsivity of hypothalamic neurons to insulin and leptin is impaired during pregnancy in rodents, although the specific cell types exhibiting this impairment are unknown.^62, 67–73^ Consistent with impaired leptin and insulin responsivity during pregnancy, we find reduced expression of several genes involved in downstream leptin and insulin signaling in AgRP and POMC neurons during pregnancy, and increased expression of the transcription factor Foxo1 in AgRP and POMC neurons during pregnancy (**Fig. 7E, F, J, and K)**. Given that Foxo1 directly inhibits the downstream effects of insulin and leptin on AgRP and POMC expression,^41, 74^ this finding provides a putative mechanism mediating impaired sensitivity of AgRP and POMC neurons to leptin and insulin and altered mRNA expression of these peptides during pregnancy. Ultimately, these changes would be expected to elevate AgRP neuron activity and reduce POMC neuron activity, although further studies are required to confirm this hypothesis. Further work is also required to map the effects of pregnancy-related neuroendocrine changes on AgRP and POMC neuron activity during pregnancy,^49^ and to determine the neurophysiological mechanisms mediating increased AgRP neuron activity and reduced POMC neuron activity during pregnancy.

Notably, for all arcuate cell types, the transcriptional changes associated with pregnancy far exceeded those associated with fasting (**Fig. 6E, H, and K**). Given the profound effect of fasting on the transcriptional state of arcuate nucleus neurons,^40, 75, 76^ the difference in magnitude between the transcriptional changes associated with fasting and pregnancy are surprising. While we cannot completely exclude that this difference in magnitude is in part due to technical effects, we were able to detect many transcriptional changes in AgRP neurons that are known to occur in response to fasting (**Fig. 6H-J**). Further, our data demonstrating an approximately 5-10 fold increase in transcriptional changes in AgRP neurons vs POMC neurons in response to fasting is consistent with prior reports utilizing cell type specific transcriptomic profiling of these neurons in fed and fasted mice.^40^ Thus, while the spatial transcriptomic approach presented here may differ in sensitivity between previously published RNA sequencing datasets on AgRP and POMC neurons, the changes we observed in response to fasting are consistent with prior reports (**Fig. 6H**). We instead suggest that the magnitude of the transcriptomic changes in arcuate neurons reported here during pregnancy vs fasting may reflect a larger magnitude of changes associated with the long-term neuroendocrine and metabolic changes associated with pregnancy, compared with the acute changes associated with fasting. Further work is required to precisely characterize the magnitude of the transcriptional changes associated with pregnancy, and the effect of these changes on neural circuit activity. However, the data presented here clearly demonstrate a drastic effect of pregnancy on the transcriptome of the arcuate nucleus and should provide a valuable resource for researchers interested in the effects of pregnancy on neuronal physiology. Further, we establish bi-directional changes in AgRP and POMC neuron activity as critical cellular nodes mediating the effects of pregnancy on energy homeostasis.

## Methods

### Animals

All experiments were approved by the University of Illinois Institutional Animal Care and Use Committee (IACUC). Experiments were performed on female mice (8-16 weeks old). Experiments were performed on C57BL6J (Jax#000664), AgRP-Cre (Jax#012899), and POMC-Cre mice (Jax# 005965). Mice for RNAscope and Xenium analysis were C57BL6J mice that were ordered from Jackson Labs. AgRP-Cre and POMC-Cre mice were bred in house by breeding Cre heterozygous mice with C57BL6J mice (which were ordered from Jackson Labs). Litters were genotyped in house with standard PCR primers for the Cre gene to confirm the transgenic allele. Primers that were used for both AgRP-Cre and POMC-Cre are Cre common (5’ GCT TCT TCA ATG CCT TTT GC 3’) and Cre mutant (5’ AGG AAC TGC TTC CTT CAC GA 3’). Prior to experiments mice were group housed in 2-5 mice per cage. Mice were housed in a temperature (20 C) and humidity-controlled environment, with a 12 h light/dark cycle. All mice had *ad libitum* access to food and water unless otherwise noted in the text. To generate pregnant and control mice for all studies female mice were bred to a reproductively experienced male mouse for 5 days. 5 days was chosen as this period covers the entire length of the mouse estrus cycle and was found in preliminary studies to result in the largest percentage of successful pregnancies. Mice were checked daily for a vaginal plug indicating successful mating. Control virgin mice were instead paired with a female mouse for 5 days. Following mating, all mice were single caged and daily food intake and body weight was measured. In animals subjected to surgical procedures (i.e. chemogenetic and fiber photometry assays) animal breeding occurred between 2-4 weeks following viral injections. All experiments were performed during the third trimester (day 14-20) of pregnancy or the equivalent time-period in non-pregnant control mice during diestrus.

### Viral Vectors

Adeno-associated viral vectors (AAV) that were used in this study included Cre-dependent GCAMP6s (AAV5-Syn-Flex-GCAMP6s-WPRE-SV40), Cre-dependent hM4Di (AAV5-hsyn-DIO-hM4Di-mCherry), Cre-dependent hM3Dq (AAV5-hsyn-DIO-hM3Dq-mCherry), and Cre-dependent mCherry control virus (AAV5-hsyn-DIO-mCherry). All viruses were purchased from addgene and were injected into the brain at stock concentrations (> 1×10^12 vg/mL).

### Pharmacology

The following pharmacological drugs and doses/routes of administration were used in this study: peptide YY (3-36) (10ug/kg, i.p.; Tocris: 1618), liraglutide (100ug/kg, s.c., Tocris: 6517), GDF-15 (10ug/kg, i.p., R&D Systems: 8944-GD), Clozapine-N-oxide (CNO, 1mg/kg, i.p.; Enzo Life Sciences: BML-NS105), Setmelanotide (100ug/kg or 300ug/kg, i.p.; 0.2ug, i.c.v.; AdooQ: A20689)

### Stereotaxic viral injections and fiber optic implants

Stereotaxic surgeries were performed as described in our prior studies. For surgeries, mice were anesthesized with isoflurane and positioned into a stereotaxic apparatus (Kopf) with a constant flow of oxygen and isoflurane. Mice were administered preoperative carprofen (5mg/kg, s.c.), and a small incision was made on the skull in the area between bregma and lambda markers. AAV vectors were injected into the arcuate nucleus using a pulled glass micropipette, which was attached to a micromanipulator (Ronal Tool). Viral injection coordinates for targeting the arcuate nucleus in AgRP and POMC-Cre mice were as follows (from bregma): A/P: −1.4 and −1.8mm, M/L: +/-0.3mm, D/V: −5.65mm and −5.8mm (from the surface of the brain). Two injections (200ul/injection) were made at each of the two A/P coordinates on both sides for chemogenetic experiments, and on one side for fiber photometry experiments. For each injection, virus was injected over 5 minutes and left for an additional 5 minutes before removing the needle to reduce viral leakage outside of the intended target site.

For fiber photometry experiments, during the same surgery, a fiber optic cannula (200um, RWD Biosciences) was implanted directly about the arcuate viral injection site at the following coordinates: A/P: −1.60, M/L: 0.2mm, D/V: −5.60 from the surface of the brain. After fiber insertion, the fiber was secured to the skull using dental cement (C&B Metabond). Following chemgenetic and fiber photometry surgeries, mice were single caged and returned to housing facilities. Animals were monitored daily for 10 days following surgeries, and behavioral experiments did not start until 3 weeks following the completion of surgeries (to allow for surgical healing and viral expression).

### Fiber photometry experiments

Fiber photometry equipment and analysis was performed as described in our prior study.^77^ For fiber photometry experiments mice were tethered to a Plexon Multi-Wavelength Fiber Photometry System (Plexon, 8-61-A-07-A) via a multi-fiber patch cord (Plexon, 08-60-A-04-C). Patch cords were directly attached to the fiber optic implant on the mouse’s head via a ceremic mating sleeve (Plexon). Blue (465nm) and UV (410nm) light sources were provided by internal LED drives connected to the fiber photometry system. The system cycles on and off at 30hz sampling window between the 410nm (isosbestic control signal) and 465nm (GCAMP6 signal). Signals are collected via an installed fluorescent camera in the Plexon system. Fiber photometry analysis was performed using a custom R code, as described in our prior study.^77^

After allowing 3 weeks for viral expression and recovery from surgery, all mice were fasted overnight and provided with a food pellet. Only animals with the expected inhibitory (AgRP) or excitatory (POMC) response to chow were included for additional experiments. Following the screening of all mice we approximately divided mice into “pregnant” and “non-pregnant” groups, with the two groups consisting of mice with an equivalent baseline calcium response to the presentation of food. The pregnant group was bred to reproductively experienced male mice following baseline experiments, while the non-pregnant group was paired with a female mouse. All mice were subsequently tested in two behavioral assays, which were performed both prior to pregnancy (baseline period) and during the third trimester of pregnancy (or the equivalent time-period in non-pregnant control mice). First, to test the response to food consumption in the hungry state, mice were sequentially fasted for 45 minutes and six hours during the dark period (ZT12-ZT18). For each recording session, mice were attached to a fiber optic patch cord and were allowed to move freely in their home-cage for 5 minutes before starting the recording. Baseline calcium signal was subsequently recorded for five minutes, at which point a “caged” food pellet was dropped in the middle of the cage. Calcium signals were measured for an additional 5 minutes following the addition of “caged” food and the change in calcium signal was determined during this period, compared to the five-minute baseline period. After 5 additional minutes, the cage was removed from the food, allowing mice to access and consume the food pellet for 5 additional minutes. Two to three days later the same mice were tested in a second assay to measure the response to peanut butter (PB) chip consumption in the *ad libitum* fed state. Mice were provided with a peanut butter chip daily in their home cage for two days prior to testing to habituate the mice to the chip and to train the mice to eat the PB chip upon presentation. For PB chip testing, mice were connected to the photometry system as previously described and baseline calcium responses were recorded for five minutes. After five minutes mice were provided with a “caged” peanut butter chip for five additional minutes, and the change in signal after PB chip presentation was compared to the five-minute baseline period. After 5 additional minutes of testing, the “cage” was removed from the PB chip and mice were allowed to consume the PB chip for 5 additional minutes. Following initial 6-hour fasting and PB chip experiments (prior to pregnancy), all mice were either mated to a male mouse (pregnancy group) or paired with a female mouse for five days (non-pregnant group). 6-hour fasting and PB chip consumption experiments were subsequently repeated on the same mice during the third trimester of pregnancy or the equivalent time-period in non-pregnant mice. The change in signal was compared between the same mice during the baseline period and during the pregnancy period for each mouse for both assays for statistical analysis (i.e. within subject’s comparisons in the same mice, Fig. 3, see figure legends for additional details). We also compared the change in calcium response in non-pregnant and pregnant mice during the same testing session (i.e. between subjects comparison, see Extended Data Fig. 2).

### Feeding behavioral assays and experiments with FED3 devices

All feeding experiments were performed with feeding experimental devices, except for pharmacological experiments (**Fig. 5 and Ext Data Fig. 3**), which were performed by manual measurements. For manual measurements of food intake mice were single housed at least one week prior to starting studies. Experiments were performed at the start of the rodent dark period (ZT12). Cages were changed daily during testing to prevent crumbs. First, all mice were injected daily with saline (i.p., 200ul) for at least 2 consecutive days to habituate to injections. After habituation, non-pregnant and pregnant mice were injected with either saline or drugs (PYY, liraglutide, GDF15) in a randomized manner. Immediately following injections, a pre-measured amount of fresh food (between 15-20 grams) was provided to the mice on their food hopper. Food intake was manually measured 2 hours following injections, and food intake was compared between saline and drug injection conditions for each mouse.

FED3 feeding assays were performed as described in our prior study.^77^ Briefly, to measure feeding structure in undisturbed mice in a home-cage environment, FED3 devices were attached to the side of the mouse’s cage. Mice obtained all their food from FED3 devices during testing. The FED3 devices were set at Fixed Ratio 1 (FR1) schedule of reinforcement in which one nose poke on the left poke, results in dispensing of one 20mg food pellet. This feeding schedule was chosen over free access feeding as we observed significantly less food hoarding in FR1 schedule vs free feeding model. The removal of the food pellet is sensed by the FED3 device, allowing the quantification of feeding patterns. Feeding patterns were characterized by meal size and meal frequency. A meal was defined as the number of pellets taken where the inter-pellet interval between each of the pellets was less than 5 minutes. The meal frequency was the number of meals taken in a 24h period. After 1-2 days of initial training, when mice had reached at least 70% correct nose pokes (i.e. 70% of nose pokes occur on the correct left poke vs the incorrect right poke), we began performing feeding studies. Data was collected every day at ZT6.

For chemogenetic feeding experiments with the FED3 devices (**Fig. 4**), mice were administered saline (i.p., 200ul) or CNO (1mg/kg, i.p.) in a randomized fashion at the start of the rodent dark period (ZT12). Changes in pellets consumed, meal size, and meal number were calculated for each mouse following saline or CNO injections and compared for statistical analysis (repeated measures within sample comparison).

### ICV administration of setmelanotide

To test the effect of melanocortin receptor stimulation directly in the central nervous system we performed a stereotaxic surgery to implant an infusion cannula into the lateral ventricle in adult female mice. Surgeries were performed as described above for virus injection and fiber optic implants except an infusion cannula (RWD Biosciences) was instead implanted into the lateral ventricle and abhorred to the skull using dental cement. A dummy cannula was attached to the infusion cannula to prevent the device from clogging. Following surgeries, mice were single caged and returned to animal housing facilities for at least one week before starting experiments. Following surgical recovery, mice were mated (or paired with a second female) to generate experimental mice as described above. Once mice reached the second trimester of pregnancy (or the equivalent time-period in non-pregnant control mice), mice were first administered control artificial cerebral spinal fluid (aCSF) for 2-3 days (1ul) to habituate to handling and i.c.v. injections. Food intake was measured manually two hours following infusions, as previously described. Next, mice were randomly administered either aCSF (1ul) or setmelanotide (0.2ug, i.c.v in 1ul aCSF) and food intake was measured 2 hours later.

### Perfusion and sectioning

Following experiments all mice were perfused with 4% paraformaldehyde followed by dissection of the brain. The brains were transferred to a fresh 4% paraformaldehyde solution for 24 hours, followed by 24 hours in 10%, 20% and 30% sucrose solutions in 1x PBS for Immunohistochemistry, RNAscope, and Xenium experiments. Brain slices containing the hypothalamus were then obtained by sectioning on a cryostat (Leica, CM3050S) at 40µm thickness (for IHC), 20um for RNAscope experiments, and 10um for Xenium experiments. For IHC and post hoc histological verification of viral injections/fiber implants, these sections were then placed into 24 well plates containing 500µL of 1x ultra-pure PBS, mounted on Superfrost glass slides (Fisher), and imaged with confocal microscopy (Zeiss LSM 700 microscope, Z-stack and tile scan of whole hypothalamus). For RNAscope experiments, sections (20µm) were mounted directly onto Superfrost microscope glass slides (Fisher scientific) and RNAscope was performed as described in “*RNAscope in situ hybridization and mRNA quantification”* described below.

### RNAscope in situ hybridization

RNAscope analysis was performed on C57/BL6J WT non-pregnant and pregnant mice that were perfused and the brain sections obtained as described above. RNAscope multiplex fluorescent in situ hybridization version 2 was used according to the protocol described in the kit. AgRP, POMC, MC4R, CRH, TRH and fos probes utilized were Mm-Agrp-C1 (Ref: 400711), Mm-Pomc-C2 (Ref: 314081-C2), Mm-Mc4r-C2 (Ref: 319181-C3), MmCRH-C1 (Ref: 316091), Mm-TRH-C1 (Ref: 436811), and Mm-Fos-C3 (Ref: 316921-C3) respectively. Following RNAscope protocols, confocal images were taken using LSM900 microscope and Zen software (z-stack and 20x zoom) and the cells were counted in the entire ARC or PVN regions using Fiji/ImageJ software. The colocalization cell count was performed considering the nuclei (stained with DAPI) containing at least two transcripts of each probe as a positive cell. For the mRNA signal intensity measurements, using Fiji/Image J, a ROI was created covering the entire ARC region and the signal was measured using the “measure” software feature. We analyzed at least 4 sections per mouse and the average per animal was used for statistical analysis, for both cell count and signal intensity comparisons.

### Immunohistochemistry and cfos quantification

For the chemogenetic experiments, the 40um brain sections were placed into 24 well plates containing 500µL of blocking buffer (100 mL Ultrapure 1X PBS, 2 grams Bovine Serum Albumin, and 100µL of Tween 20) and placed on a shaker at room temperature and allowed to sit for 2 hours. Next, a master mix of the rabbit cFos (1:1000; 9F6 Rabbit mAb, Cell Signaling Technology) primary antibody in blocking buffer was prepared. 500µL of this master mix was added to each well-containing brain sections and placed onto a shaker at 4°C overnight. The primary antibody mixture was replaced by 500µL of ultra-pure 1X phosphate-buffered saline (PBS) and placed on a shaker at room temperature for 10 minutes and this step was repeated twice more. The secondary antibody (Goat anti-Rabbit IgG (H+L) Cross-Adsorbed Secondary Antibody, Alexa Fluor™ 488) was prepared in blocking buffer in a 1:500 concentration and then 500µL of the secondary was added to each well and placed on a shaker, at room temperature, and incubated for 2 hours. Following three 10-minute wash steps with 1X ultra-pure PBS, the sections were mounted on Superfrost glass slides (Fisher scientific) and analysis of the images were performed using LSM900 confocal microscope (Z-stack, 20x).

At least 3 sections with mcherry expression in the arcuate nucleus per mouse were used to count the mcherry and cfos signals, and the average of those measurements was calculated for each mouse and used in the statistical analysis. The cfos quantification was performed using ImageJ/Fiji software and the total number of AgRP or POMC cells expressing fos was counted and the percentage of these cells co-expressing with cfos was quantified for the statistical analysis.

### Xenium spatial transcriptomics preparation

Diestrus non-pregnant ad libitum, diestrus non-pregnant 24h fasted, and third-trimester pregnant ad libitum female mice for the xenium experiment were perfused and the brains treated with sucrose solutions as described in “*<u>Perfusion and sectioning</u>*”. The brains were then sliced using a 1 mm Coronal Mouse Brain Slicer with razor blades at most anterior and most posterior hypothalamus (A/P: −1.00mm∼ −2.30mm) and the medial hypothalamus was quickly dissected from each coronal section and immediately placed in the container (see Figure 6). After the dissection and placement of all medial hypothalamus in the container, a solution of equal parts of OCT and 30% sucrose was used to cover the medial hypothalamus segments, and it was quickly frozen for cryostat sectioning. All the brains were then sectioned at the same time as a block until the area of interest for the anterior medial hypothalamus (A/P: −1.50mm) and the posterior medial hypothalamus (A/P: −1.90mm). Each one of the 10um medial hypothalamus sections were immediately placed on pre-frozen Xenium slides and maintained in −80°C freezer until the start of the Xenium run.

### Xenium spatial transcriptomics analysis

Slides were imaged on a Xenium analyzer machine by following the manufacturer-recommended protocol by an on-site omics facility (University of Illinois Roy J. Carver Biotechnology Center Cytometry and Microscopy to Omics Core Facility). Cell segmentation and transcription localization were compiled by the Xenium Onboard Analysis pipeline, and transcripts were assigned to cells following the Xenium defined cell segmentations software. Divisions of the medial hypothalamus corresponding to the arcuate nucleus and median eminence were extracted for each mounted section following expert annotation and used to generate cell-by-transcript matrices for each independent sample. These were then combined into a single Seurat object,^78^ for downstream clustering and annotation.

*Spatial single-cell clustering:* The combined “single-cell” data was analyzed using Seurat v5.2.1, and functions are derived from this package unless stated otherwise.^78^ The Xenium count data was normalized via SCTransform^79^ using glmGamPoi. Annotation to broad neuronal and glial cell types was achieved using a combination of supervised and unsupervised techniques. The first pass of annotation was performed using SingleR^80^, with the murine Hypomap^76^ acting as a reference dataset. Briefly, SingleR was trained on log-normalized and scaled Hypomap counts, and author annotated class labels (*e.g*. “Neurons”, “Tanycytes”, “Astrocytes”, *etc.*) to form correlative associations between transcriptional state and cell-type and thus predict labels for our Xenium data. Compared to the Hypomap, annotations of our Xenium data were enriched for tanycytes and ependymal cells, which is consistent with our anatomical location around the arcuate nucleus and 3rd ventricle (Figure 6C). Transcriptional state similarity was high between developmentally linked glial cells, *e.g.* astrocytes, tanycytes and ependymal cells, and cells of endothelial lineage. Where microanatomical location of cells determines likely cell type (*e.g.* tanycytes lining the 3rd ventricle), the agreement between SingleR predicted assignment and location was extremely high (Ext Fig. 4).

Unsupervised clustering was performed in parallel, via the standard Seurat pipeline of principal component analysis (PCA) (RunPCA()), construction of a shared nearest-neighbor graph on the first 30 PCA dimensions (FindNeighbors()), and finally cluster identification via the Leiden algorithm (FindClusters()). Unsupervised clusters were annotated with their majority SingleR-predicted label if within-cluster agreement was high (*i.e.* fewer than 5% of cells within the cluster were assigned a different label). Manual annotation via marker genes was performed on poorly annotated clusters that failed to be uniquely annotated by SingleR. These included microglia, a cell-type largely missing from the murine Hypomap, and cells of the neurovascular niche. Further refinement of the neurovascular and endothelial cells was done by finding marker genes (via the Wilcoxon Rank Sum variant of FindMarkers() implemented in Seurat)^78^ and matching to markers from the Allen Brain Transcriptomic Cell Atlas^81^.

*Neuronal Single Cell Clustering:* Cells predicted to be neurons by the combined supervised-unsupervised clustering were subset from the heterogeneous population, and re-analyzed with SCTransform, thus allowing variance modeling to focus on intra-neuronal differences. At each stage of clustering, data were reviewed for stratification by sample, pregnancy or feeding status (*i.e* animal condition), and technical aspects of spatial sequencing (*e.g.* transcript count and feature diversity). To reduce the potential of clustering by condition, the SCTransform model regressed the effect of condition via the “vars.to.regress” parameter. Unsupervised clustering resolutions were chosen by maximizing average cluster Silhouette scores using PCA embeddings and inspecting clusters for markers of canonical neuroendocrine and centrally-projecting neurons of the murine arcuate nucleus. Marker genes for clusters were found using FindMarkers() using Wilcoxon Rank Sum tests, as before.

Neurons that expressed markers that were indicative of a non-arcuate origin were mapped back to their histological origin for validation.

*DESeq2 Modeling:* While differential gene expression between cells obtained from different “treatments” is possible on a single-cell basis using an SCTransform model, this approach lacks sample-level error modeling and treats individual cells from the same tissue as independent. Gene expression changes induced by fasting or pregnancy across the cell-types of the arcuate nucleus were instead modeled at the tissue sample level using DESeq2 (v1.40.2).^82^ Raw count data was “pseudo-bulked” by summing total counts across cells by sample (*i.e.* transcripts from tissue obtained from a single animal) and split by assigned cell cluster. By incorporating slide origin as a batch covariate and specifying variation from *ad-libitum* as contrasts of interest, the change in expression attributable to condition was modeled separately from technical covariates of sample depth, and batch variation. DESeq2 fits a mean-dispersion trend that utilizes global structure of expression data and so combining counts from *ad-libitum* control mice, 24-hour fasted mice, and pregnant mice has a tendency to shrink the moderate transcriptional changes induced by fasting, in favor of the greater transcriptional variation caused by pregnancy. To account for this, separate DESeq2 models were fit for *ad-libitum* vs. fasted and *ad-libitum* vs pregnant individually, alongside a joint multi-level model for global comparisons for each neuron and glial cell type. Samples were excluded from the cell-type DESeq2 model if they failed either of the following thresholds: the sample contained fewer than 15 cells reliably mapped to cell-type of interest, and/or correlation of the sample with respect to its anterior/posterior opposite was very low. In practice, this procedure excluded tissue that exhibited histological damage or was too anterior to capture the arcuate nucleus.

*Differential gene expression & sample-wide expression:* Differential gene expression results are reported for *ad-libitum* vs. fasted and *ad-libitum* vs pregnant using the split models, and utilize the “Approximate Posterior Estimation for generalized linear model” (apeglm)^83^ log-fold change shrinkage procedure to improve fold-change estimation for low count genes. Sample-level expression heatmaps (fig. 6M, 7F, 7Q-S) use counts corrected by DESeq2 estimated size factor, and scaled genewise via simple min-max normalization to a range of 0-10 for plotting. To enable comparisons between Pomc and AgRP neurons, corrected counts from AgRP and Pomc neurons were combined prior to scaling. Between-sample correlation heatmaps (*fig. 7D* and 7O), and PCA plots, (*fig. 7B* and 7M), were generated using cor() and prcomp() (base R, v4.3.2) on regularized log-transformed counts via rlog() of the global DESeq2 model. The Gene-loading analysis (*fig. 7C* and 7N) extracts the Data are plotted using ComplexHeatmap,^84^ EnhancedVolcano^85^, and ggplot2^86^ for heatmaps, volcano-plots and other figures respectively.

*Xenium per cell transcription analysis:* To analyze the transcription of individual AgRP or POMC cells, we utilized the Xenium Explorer 3 software. Cells with the mRNAs of interested required at least 2 transcripts to be considered positive.

### Statistical Analysis

Data that was normally distributed was analyzed with parametric tests, while data that was not normally distributed was analyzed with non-parametric tests. All statistical tests and statistics are detailed in the respective figure legends. Data was either analyzed with Graphpad Prism or R (Xenium data analysis).

---

**Extended Figure 1:**
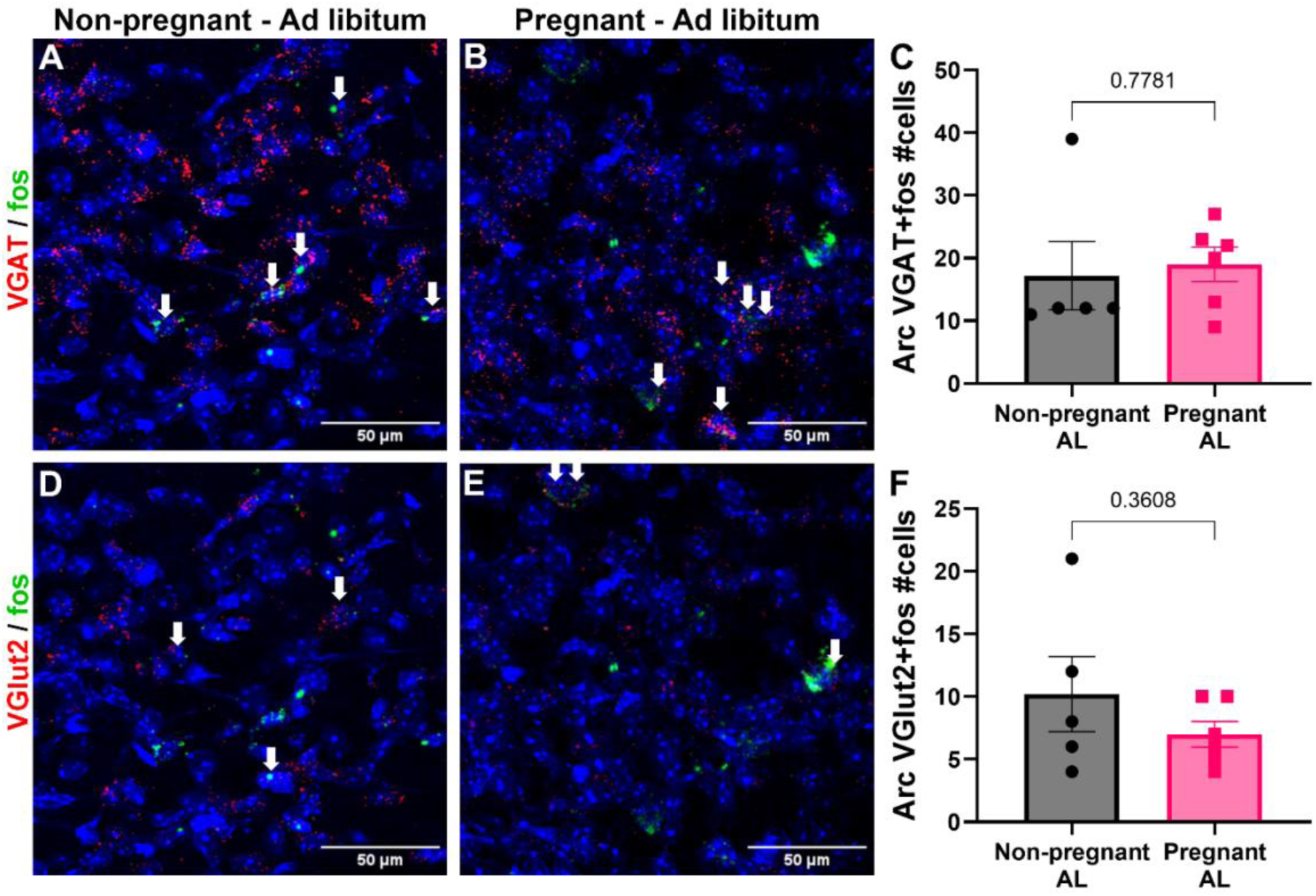
Pregnancy does not change the activity of VGAT or VGlut2 expressing cells in arcuate nucleus. (A-B) Confocal images of VGAT (red) and fos (green) mRNA expression and DAPI (blue) in the Arc of non-pregnant (A) and pregnant (B) mice with *ad libitum* access to food (L). Comparison between the number of VGAT expressing cells colocalized with fos in non-pregnant and pregnant mice with *ad libitum* access to food (n=5-6). (D-E) Confocal images of VGlut2 (red) and fos (green) mRNA expression and DAPI (blue) in the Arc of non-pregnant (D) and pregnant (E) mice with *ad libitum* access to food (L). Comparison between the number of Vglut2 expressing cells colocalized with fos in non-pregnant and pregnant mice with *ad libitum* access to food (n=5-6). Data are presented as mean values ± SEM. For all panels, statistical significance was tested by unpaired t-test. For all panels *p\** < 0.05, *p\*\** < 0.01, *p\*\** < 0.001. Data points represent individual mice.

**Extended Data Fig. 2:**
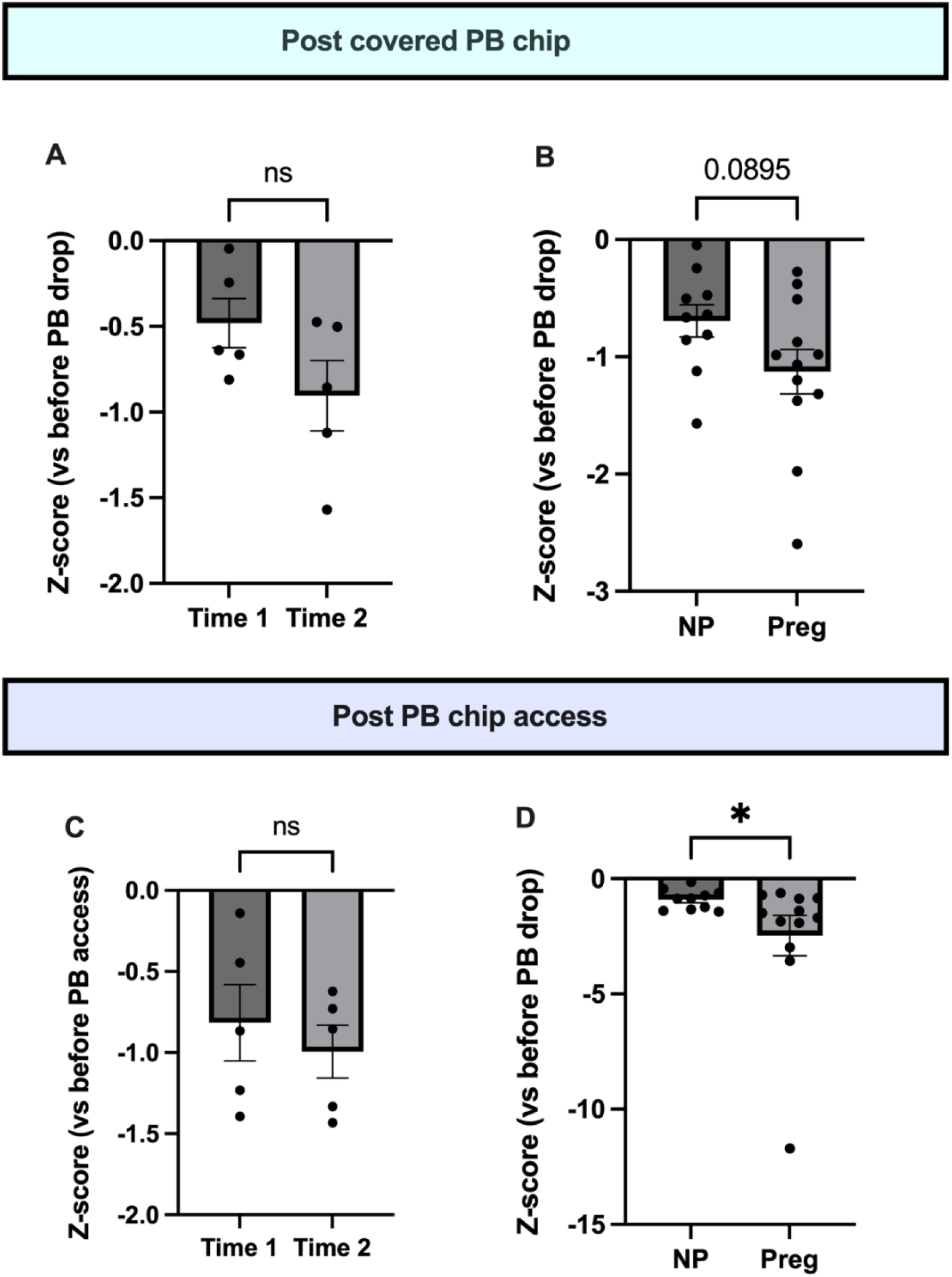
Pregnancy increases inhibitory response to PB chip presentation. (A) Calcium response to covered PB chip during initial recording and during time-matched recording during the pregnancy period in non-pregnant control mice. (B) Change in calcium activity in non-pregnant and pregnant mice (between subjects comparison) during the pregnancy period. (C) Calcium response to PB chip access during initial recording and during time-matched recording during the pregnancy period in non-pregnant control mice. (D) Change in calcium activity in non-pregnant and pregnant mice (between subjects comparison) during the pregnancy period. Data points represent individual mice. Panels A and C analyzed by Wilcoxon matched-pairs signed rank test, panels B and D analyzed by Mann Whitney test.

**Extended Data Fig. 3:**
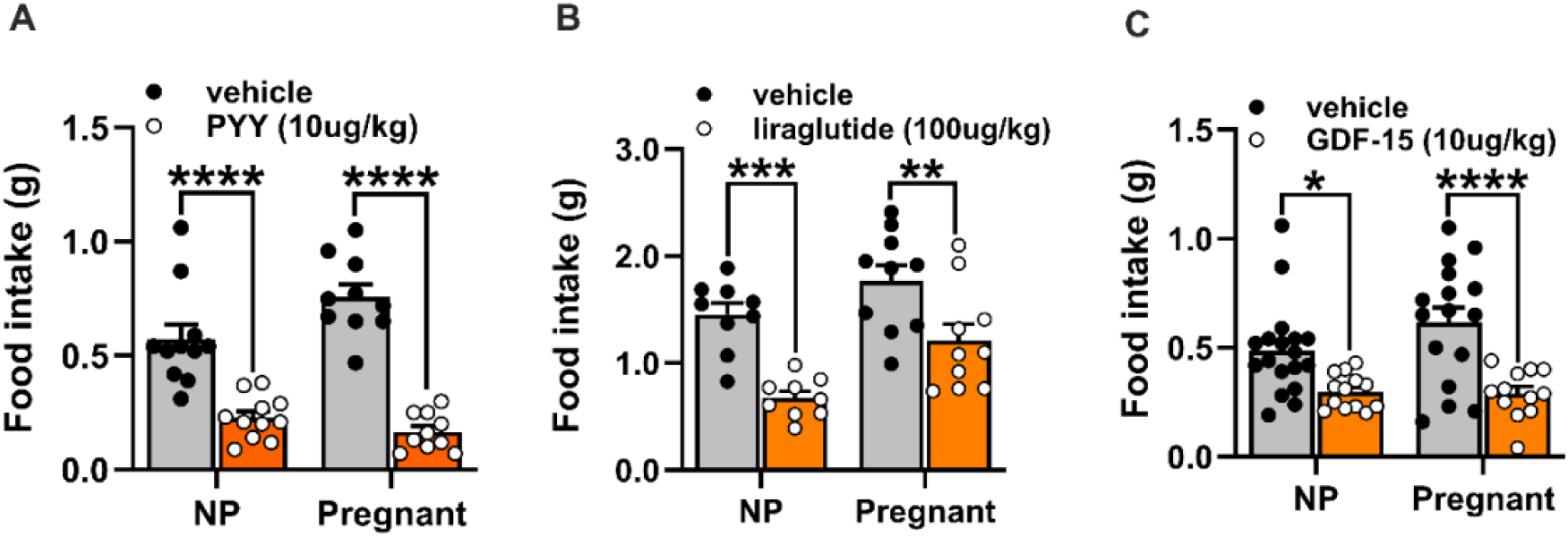
Pregnancy does not alter anorexic response to PYY, liraglutide, or GDF-15. Food intake in non-pregnant (NP) and pregnant mice following systemic injection of PYY (A), the GLP1-R agonist liraglutide (B), and GDF-15 (C). Data points represent individual mice. Data analyzed by Two-way ANOVA with Sidak post hoc test: *, p<0.05; **, p<0.01; ***, p<0.001; ****, p<0.0001.

**Extended Figure 4:**
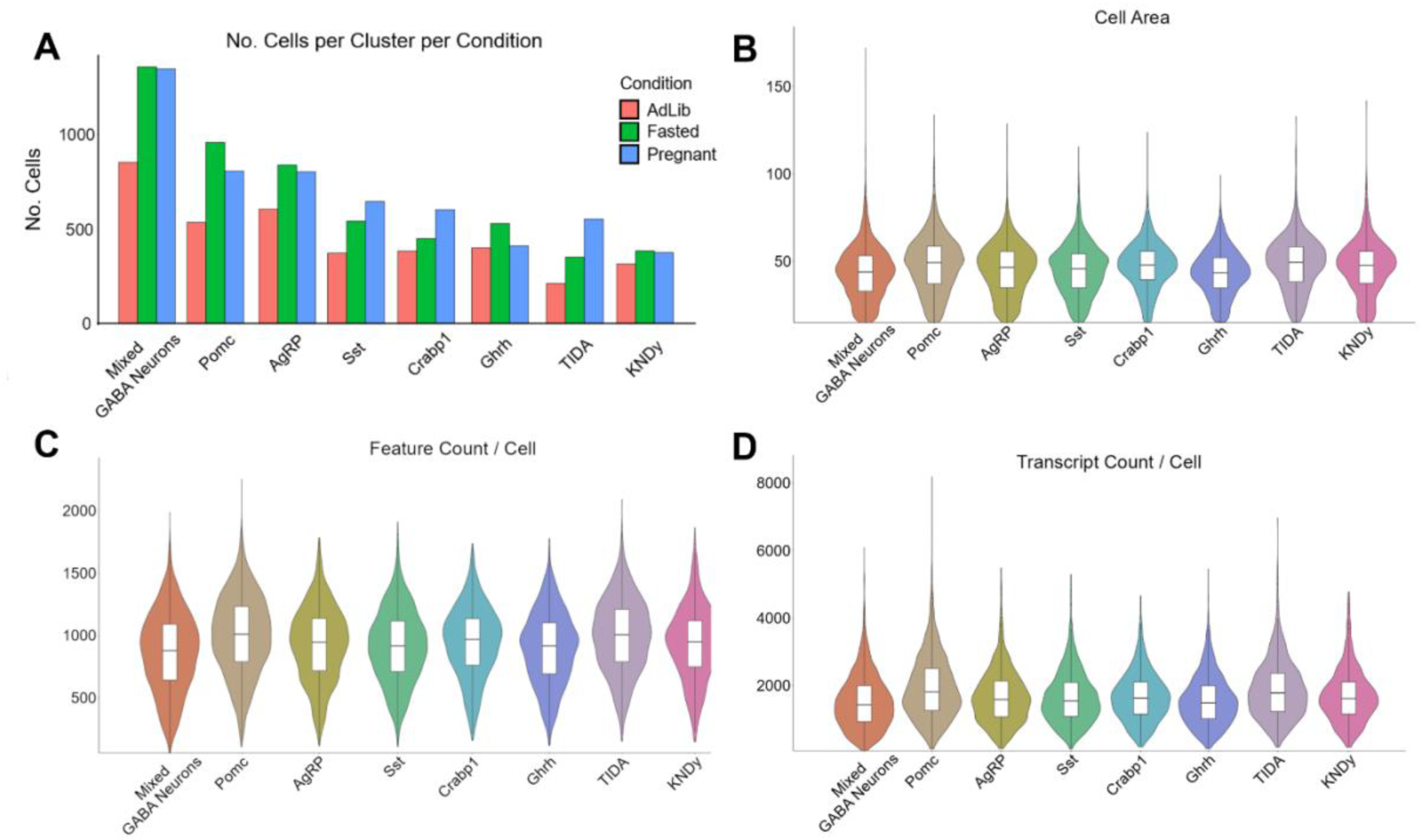
Quality control data for arcuate Xenium analysis. (A) Number of cells present in different neuronal clusters per condition group: non-pregnant *ad libitum* females (red), non-pregnant 24-hour fasted females (green), and pregnant *ad libitim* (blue) (B-D) Violin plots of all experimental groups combined and their neuronal clusters individual cell area (B), feature count per cell (C), and transcript count per cell (D). For all panels n=4-5 mice per group.

